# A Unifying Mechanism Governing Inter-Brain Neural Relationship During Social Interactions

**DOI:** 10.1101/2021.06.02.446694

**Authors:** Wujie Zhang, Michael M. Yartsev

**Affiliations:** Helen Wills Neuroscience Institute and Department of Bioengineering, UC Berkeley, Berkeley, 94720, United States

## Abstract

A key goal of social neuroscience is to understand the relationship between the neural activity of socially interacting individuals. Decades of research have focused on a single aspect of that relationship: the similarity in neural activity across brains. Here we instead asked how neural activity differs between brains, and how that difference evolves alongside activity patterns shared between brains. Applying this framework to pairs of bats engaged in spontaneous social interactions revealed two complementary phenomena characterizing the inter-brain neural relationship: fast “inter-brain catch-up” unfolding in parallel with slow activity covariation across brains. A model reproduced these observations, generated multiple predictions that we confirmed using experimental data, and provided testable hypotheses for studying the inter-brain relationship in larger social groups. Together, the data and model suggest a parsimonious computational mechanism—opposite feedback to neural activity components reflecting inter-brain difference and similarity—that unifies diverse aspects of the inter-brain neural relationship.

## Introduction

What is the relationship between the neural activity of socially interacting individuals? This central question in social neuroscience has motivated nearly two decades of research spanning a diversity of species and methodologies (e.g. Babiloni and Astolfi, 2014; Dumas et al., 2011; Freiwald, 2020; Hasson et al., 2012; Hasson and Frith, 2016; Hoffmann et al., 2019; Kingsbury and Hong, 2020; Koike et al., 2015; Konvalinka and Roepstorff, 2012; Liu et al., 2018; Montague et al., 2002; Redcay and Schilbach, 2020; Scholkmann et al., 2013; Schoot et al., 2016; Testard et al., 2021; Tseng et al., 2018; Wass et al., 2020). Yet, despite this diversity, nearly all research has tackled the study of inter-brain relationship under a single conceptual framework: considering the neural activity of two interacting individuals as the two variables of interest, and searching for similarities between them. Commonly, similarity was assessed using measures related to either correlation (Dikker et al., 2014; Kawasaki et al., 2013; King-Casas et al., 2005; Kingsbury et al., 2019; Kinreich et al., 2017; Levy et al., 2017; Liu et al., 2017; Montague et al., 2002; Piazza et al., 2020; Silbert et al., 2014; Spiegelhalder et al., 2014; Stephens et al., 2010; Stolk et al., 2014; Tomlin et al., 2006; Zadbood et al., 2017; Zhang and Yartsev, 2019) or coherence (Cui et al., 2012; Dikker et al., 2017; Dumas et al., 2010; Goldstein et al., 2018; Levy et al., 2017; Lindenberger et al., 2009; Montague et al., 2002; Mu et al., 2017; Stolk et al., 2014; Yang et al., 2020; Yun et al., 2012). The extensive research under this framework has provided a detailed description of the phenomena of inter-brain correlation and coherence across a wide variety of conditions.

However, one aspect of the inter-brain neural relationship has remained unexplored: the difference in neural activity between brains. We hypothesized that inter-brain difference is more than simply a lack of inter-brain correlation, and the detailed dynamics of the difference between brains can be as informative as the similarity for understanding the inter-brain relationship. Therefore, here we took an approach that focused on both the similarity and difference in inter-brain activity, and on how the two co-evolve over time. We applied this new framework to neural activity simultaneously recorded from pairs of socially interacting Egyptian fruit bats (*Rousettus aegyptiacus*), a mammalian species known for its high level of sociality (Herzig-Straschil and Robinson, 1978; Harten et al., 2018; Prat et al., 2015, 2016, 2017; Omer et al., 2018; Cvikel et al., 2015; Egert-Berg et al., 2018; Kwiecinski and Griffiths, 1999).

As detailed below, we began by decomposing the neural activity of pairs of bats into two components representing inter-brain difference and similarity respectively. Exploratory analyses revealed that the relative timescales and variances of the two components are robust neural signatures of social interaction; moreover, their relative timescales is independent from the phenomenon of inter-brain correlation, while their relative variances is related to inter-brain correlation. Next, in search of the computational mechanism behind these findings and inter-brain correlation, we developed a model that parsimoniously explains our present observations as well as previously reported observations regarding the inter-brain relationship, uncovering a simple mechanism involving positive and negative feedback. Furthermore, the model generated multiple experimentally testable predictions regarding the inter-brain relationship. Turning back to the experimental data, we then tested and confirmed these predictions, further validating the model. Finally, we extended the model to social interactions involving more than two individuals, providing a framework for studying inter-brain relationship in larger social groups, since such group interactions often occur in real life. Combined, the insights from the experimental and modeling approaches support a unifying computational mechanism that governs the inter-brain neural relationship during social interactions.

## Results

### Relative magnitudes and timescales of inter-brain difference and mean components

We performed wireless extracellular neural recording simultaneously from pairs of bats, targeting their frontal cortex, a region implicated in social cognition in rodents, bats, non-human primates, and humans (Adolphs, 2001; Amodio and Frith, 2006; Cao et al., 2018; Chang et al., 2013; Eliades and Miller, 2017; Forbes and Grafman, 2010; Haroush and Williams, 2015; Kingsbury et al., 2019; Liang et al., 2018; Miller et al., 2015; Nummela et al., 2017; Ong et al., 2020; Pearson et al., 2014; Rudebeck et al., 2008; Tremblay et al., 2017; Zhang and Yartsev, 2019; Zhou et al., 2017). From the recordings we analyzed four types of neural signals: local field potential (LFP) power in the 30-150 Hz and 1-29 Hz bands (previously identified as relevant frequency bands in bat frontal cortical LFP [Zhang and Yartsev, 2019]; see also Supplementary Information section 3.9), multiunit activity, and single unit activity (Supplementary Information sections 3.6-3.8). We first examine 30-150 Hz LFP power, the neural signal which has been shown to have the strongest inter-brain correlation (Zhang and Yartsev, 2019), and then turn to the other neural signals.

In the experiment, pairs of bats behaved freely and interacted with each other inside a chamber (“one-chamber sessions”; Figure 1A). Under the traditional framework for studying the inter-brain relationship, one would plot the neural activity of each of the bats and see a high degree of similarity between brains (Figure 1B), as demonstrated previously (Zhang and Yartsev, 2019). This framework highlights the salience of inter-brain similarity, but makes it easy to overlook the detailed dynamics of the difference between brains. We therefore sought an analysis framework that would enable us to explicitly examine inter-brain difference and similarity side by side.

**Figure 1.**
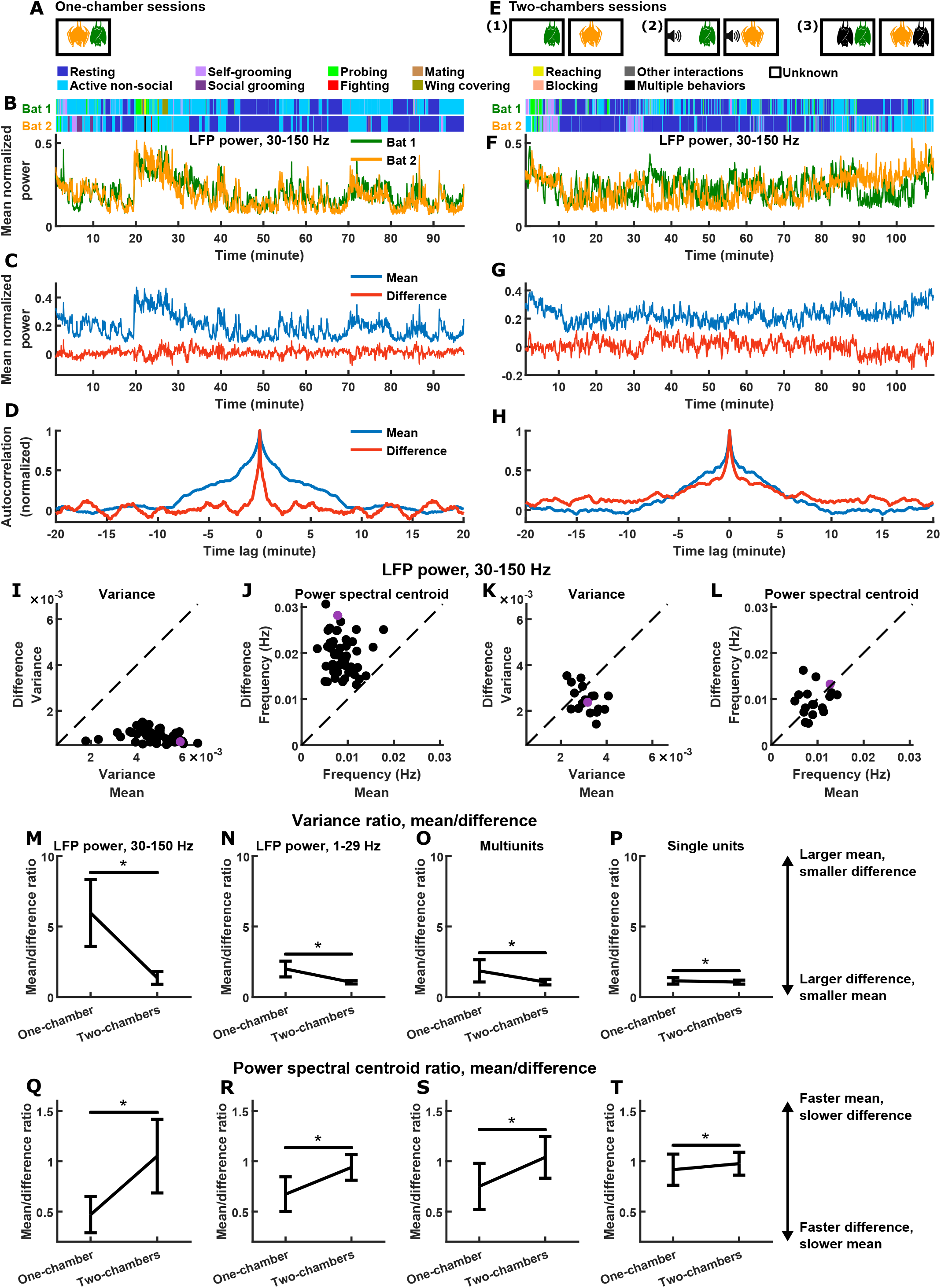
Relative magnitudes and timescales of the inter-brain difference and mean components. (A) In each one-chamber session, two bats freely interacted with each other while neural activity was wirelessly recorded simultaneously from their frontal cortices. (B) Mean normalized LFP power in the 30-150 Hz band (Supplementary Information sections 3.7, 3.9), averaged across all channels for each bat, on an example one-chamber session. Shown above are the behaviors of the two bats as a function of time, which were manually annotated frame-by-frame from recorded video. (C) The neural activity of the two bats from (B) after a change of basis, showing the mean and difference between bats. At a given time *t*, the mean and difference components are defined as 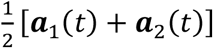 and 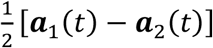, respectively, where ***a***_1_ (*t*) and ***a***_2_ (*t*) are respectively the neural activity of bat 1 and bat 2 plotted in (B). Note that the mean component had a large variance, whereas the difference component had a small variance, hovering around zero. (D) Autocorrelations (peak-normalized) of the mean and difference components shown in (C). The autocorrelations were computed after subtracting from each time series its average over time. Note that the difference component varied on faster timescales than the mean component. (E) In each two-chambers session, the same bats from the one-chamber sessions freely behaved in separate, identical chambers, while neural activity continued to be simultaneously and wirelessly recorded from their frontal cortices. There were three types of two-chambers sessions (Supplementary Information sections 3.1): (1) two bats each freely behaving in isolation; (2) two bats each freely behaving while listening to identical auditory stimuli (playback of bat calls); (3) two bats each freely behaving and interacting with a different partner in separate chambers. (F)-(H) Same as (B)-(D), respectively, but for an example two-chambers session (type 3 from (E)). Note that variances and timescales are more comparable between the mean and difference components compared to the example one-chamber session. (I)-(J) Variance (I) and power spectral centroid (J) of mean normalized 30-150 Hz LFP power, for the mean and difference components. Each dot is a single one-chamber session (the purple dot is the session shown in (B)-(D)). Variance quantifies activity magnitude, and power spectral centroid quantifies timescale (higher centroids mean faster timescales). Note that, on every one-chamber session, the difference component was smaller and faster than the mean component. The dotted lines are unity. Note that the power spectral centroid was calculated from time series of mean normalized LFP power (e.g. as plotted in (C)), not from time series of LFP itself. (K)-(L) Same as (I)-(J), but for two-chambers sessions. The purple dot is the session shown in (F)-(H). The mean and difference components have comparable magnitudes and timescales in the two-chambers sessions. (M)-(P) The average variance ratio (mean component variance divided by difference component variance) for mean normalized 30-150 Hz LFP power (M), mean normalized 1-29 Hz LFP power (N), multiunits (O), and single units (P). The averages were taken across sessions for LFP power, and across unit pairs (pooled from all sessions) for multiunits and single units. Error bars denote standard deviations. *, p<0.05, Wilcoxon rank sum test. (Q)-(T) Same as (M)-(P), but for average power spectral centroid ratio (mean component centroid divided by difference component centroid). Note that, for all four neural signals, the difference component was smaller and faster than the mean component on one-chamber sessions. See Figures S1-3 for examples and detailed results for 1-29 Hz LFP power, multiunits, and single units, respectively.

Under the traditional framework, the two variables of interest are the neural activity of each brain (e.g., in Figure 1B, each variable is the normalized LFP power averaged across the recording channels of one brain). We can represent them as a two-dimensional vector 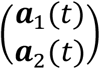, where ***a***_1_(*t*) and ***a***_2_(*t*) are the activity of bat 1 and bat 2 at time *t*, respectively. Through a change of basis, the same activity can be represented under another orthogonal basis as the mean and difference between the two brains: 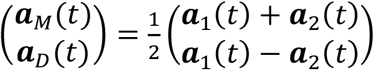, where ***a***_*M*_(*t*) is the mean component of the activity, and ***a***_*D*_(*t*) is the difference component (here the difference component is defined as 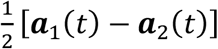 rather than ***a***_1_(*t*) − ***a***_2_(*t*) so as to have the same scale as the mean component). ***a***_*M*_(*t*) represents the common activity pattern shared between the brains, i.e., what is similar between the brains; ***a***_*D*_(*t*) represents the moment-to-moment activity difference between the brains. Note that the two components can vary independently—a large (or small) ***a***_*M*_(*t*) does not necessarily imply a small (or large) ***a***_*D*_(*t*).

Figure 1C shows the mean and difference activity components for the example session from Figure 1B. Visualizing the two components side-by-side, two relationships become immediately apparent. First, the mean component had much larger variance than the difference component (Figure 1B). This is expected given the high degree of similarity in neural activity between the brains: while the variances are not entirely determined by inter-brain correlation, a positive correlation does mathematically imply larger variance for the mean compared to the difference component (Supplementary Information section 4.3). Second, the difference component evolved over time at much faster timescales than the mean component. This is even more apparent when examining the autocorrelations of the mean and difference components (Figure 1D), where the narrower autocorrelation of the difference indicates that it varied faster than the mean.

A difference component of zero means the neural activity of the two brains are the same. The fact that the difference hovered around zero at fast timescales amounts to a phenomenon of “inter-brain catch-up”: whenever the activity of the two brains diverged from each other (i.e., the difference moved away from zero), the two quickly caught up (i.e., the difference decayed back towards zero). This rapid inter-brain catch-up process unfolded in parallel with the slow, large-magnitude activity patterns shared across the brains (i.e., the mean component), and together they characterized the inter-brain neural relationship during social interactions.

An immediate question that arises is: to what extent are these phenomena specific to social interactions? To explore potential non-social contributions to the inter-brain neural relationship, we recorded neural activity simultaneously from the same bats as they freely behaved in separate, identical chambers (“two-chambers sessions”; Figure 1E; Supplementary Information section 3.1). Normalized 30-150 Hz LFP power is shown for an example two-chambers session in Figure 1F-G. Unlike for the one-chamber session, the variances and timescales of the mean and difference components were comparable (Figure 1G-H).

We next quantified the magnitudes and timescales of the mean and difference components for 30-150 Hz LFP power on all sessions, using variance as a measure for magnitude, and power spectral centroid as a measure for timescale (Figure 1I-L; Supplementary Information section 4.2). The relationships seen in the one-chamber example above was robust: on every one-chamber session, the difference component had smaller magnitudes (Figure 1I) and faster timescales (Figure 1J) than the mean component (note that the relative magnitudes are a reflection of inter-brain correlation). On two-chambers sessions, in contrast, the mean and difference were comparable (Figure 1K-L). These relationships are summarized in Figure 1M-T for all four neural signals: LFP power in the 30-150 Hz and 1-29 Hz bands, multiunits, and single units all showed the same significant trends (see Figures S1-3 for examples and detailed results for 1-29 Hz LFP power, multiunits, and single units).

Thus, the inter-brain neural relationship during social interactions is characterized by two robust signatures of the mean and difference components: their relative magnitudes that reflect inter-brain correlation, and their relative timescales. Why is this the case? Are the relative timescales tied to the relative magnitudes? And what can these observations reveal about the mechanisms governing the inter-brain relationship? We next turn to explore possible answers to these questions.

### The relationship between relative timescales, relative magnitudes, and inter-brain correlation

In one-chamber sessions, the difference component had smaller magnitudes and faster timescales than the mean component. While the relative magnitudes of the two components are expected given inter-brain correlation (Supplementary Information section 4.3), what about their timescales? Are the observed relative timescales of the two components a necessary mathematical consequence of high inter-brain correlations, large mean components, and small difference components? Intuitively, one might think the answer would be yes: perhaps a high inter-brain correlation implies that the activity of the two brains stayed close to each other on fast timescales, i.e., implying a small difference component varying on fast timescales. This intuition is wrong. As we show in Supplementary Information (section 4.3), having a given combination of correlation, mean component variance, and difference component variance does not place constraints on the timescales of the two components. Specifically, it does not constrain the difference to be faster than the mean. To explicitly demonstrate this, we generated a set of surrogate data (Figure 2). The surrogate data was designed to have inter-brain correlation, mean component variance, and difference component variance identical to those of the actual data. However, unlike the actual data, the surrogate data was designed to have difference components that were slower than the mean components, the opposite of what was observed in the actual data (Figure 2C, E). Thus, the relative timescales of the difference and mean components are not dictated by their relative magnitudes or by inter-brain correlation. What, then, might explain the robust relationships observed between the timescales and magnitudes? And are there separate mechanisms responsible for the observed relative timescales on the one hand, and the observed relative magnitudes (and the related phenomenon of inter-brain correlation) on the other? To address these questions, we turn to computational modeling, which has emerged as a powerful tool for inferring neural mechanisms underlying social behaviors (e.g., Calhoun et al., 2019; Clemens et al., 2015). Thus, we next model the observed neural activity to infer the computational mechanisms governing the inter-brain difference and mean components, and by extension, mechanisms underlying inter-brain correlation.

**Figure 2.**
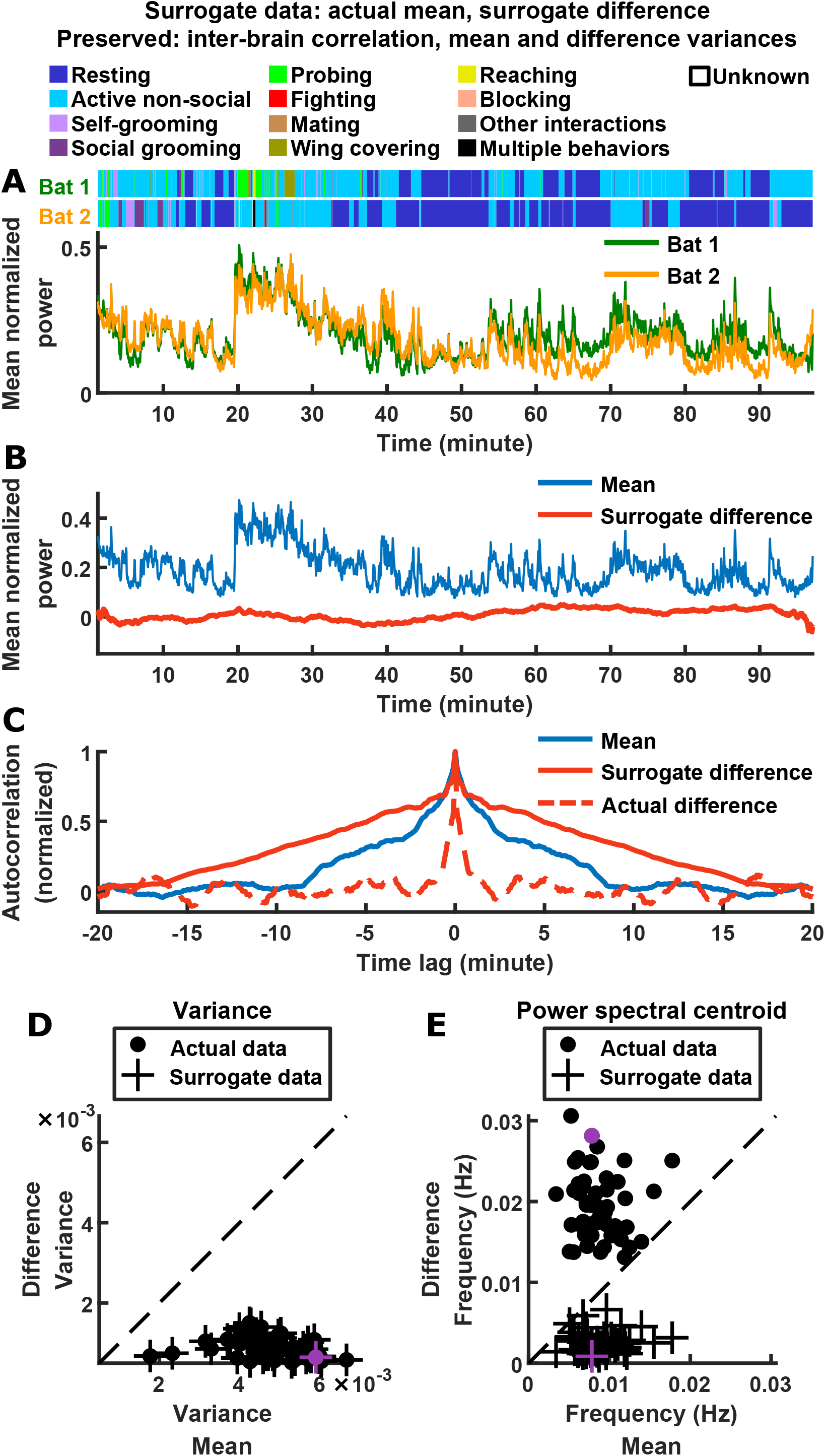
Relative timescales of the difference and mean components are not determined by their relative magnitudes or levels of inter-brain correlation. (A) Surrogate data generated from the actual data shown in Figure 1B, by combining the actual mean component with a surrogate difference component (Supplementary Information section 4.3). The surrogate was tailored such that the difference component variance and inter-brain correlation of the original experimental data were preserved (the mean component variance was also preserved since the actual mean component was used in the surrogate data). Shown above are the actual behaviors of the two bats replotted from Figure 1B. (B) The surrogate data from (A) plotted as the activity of its mean and difference components. The mean here is identical to the actual mean in Figure 1C, while the difference here has the same variance as the actual difference in Figure 1C, but with slower timescales. (C) The autocorrelations (peak-normalized) of the mean and difference components shown in (B) and of the actual difference component shown in Figure 1C. The autocorrelations were computed after subtracting from each time series its average over time. Note that for the surrogate data, the difference was slower than the mean, the opposite of what was observed experimentally. (D)-(E) Variance (D) and power spectral centroid (E) of mean normalized 30-150 Hz LFP power, for the mean and difference components of the actual data and surrogate data. Each dot is actual data from a single one-chamber session (replotted from Figure 1I-J), and each plus is surrogate data generated from the actual data of a single one-chamber session (purple dots and pluses denote the example session shown in Figure 1B-D and (A)-(C)). The dotted lines are unity. The surrogate data preserve the actual inter-brain correlations (Supplementary Information section 4.3), as well as the variances of the actual mean and difference components (pluses in (D) are at the same positions as the dots), but have slower difference components than mean components (pluses below the unity line in (E)).

### Model suggests a feedback mechanism governing inter-brain neural relationship

We modeled the neural activity of two bats using a linear differential equation (see Figure 3A for the equation; Dayan and Abbott, 2005). In the model, the neural activity variables interact through the functional coupling matrix 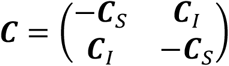, where ***C***_*S*_ is the strength of functional self-coupling and ***C***_*I*_ is the strength of functional across-brain coupling. When modeling one-chamber sessions, ***C***_*I*_ > 0, so that the activity of each bat influences the other bat’s activity. This functional across-brain coupling models the effects of sensorimotor interactions and attentional processes (Hasson et al., 2012): for example, when bat 1’s neural activity increases due to its active movements (Gervasoni et al., 2004; McGinley et al., 2015), the movements create sensory inputs to bat 2, which can drive bat 2’s neural activity to the extent that bat 2 is paying attention (Chun et al., 2011; Driver, 2001; Fritz et al., 2007; Reynolds and Chelazzi, 2004). On the other hand, when modeling two-chambers sessions, ***C***_*I*_ = 0, so that the activity of each bat does not influence the other bat’s activity. Furthermore, in both one-chamber and two-chambers models, the activity of each bat is modulated by its own behavior, which is simulated using Markov chains based on empirical behavioral transition frequencies (Supplementary Information section 4.4). Note that the usage of the term “functional across-brain coupling” in our model should be distinguished from the sense it is sometimes used in the literature to simply denote a presence of neural correlation or coherence across brains (e.g., Levy et al., 2017).

**Figure 3.**
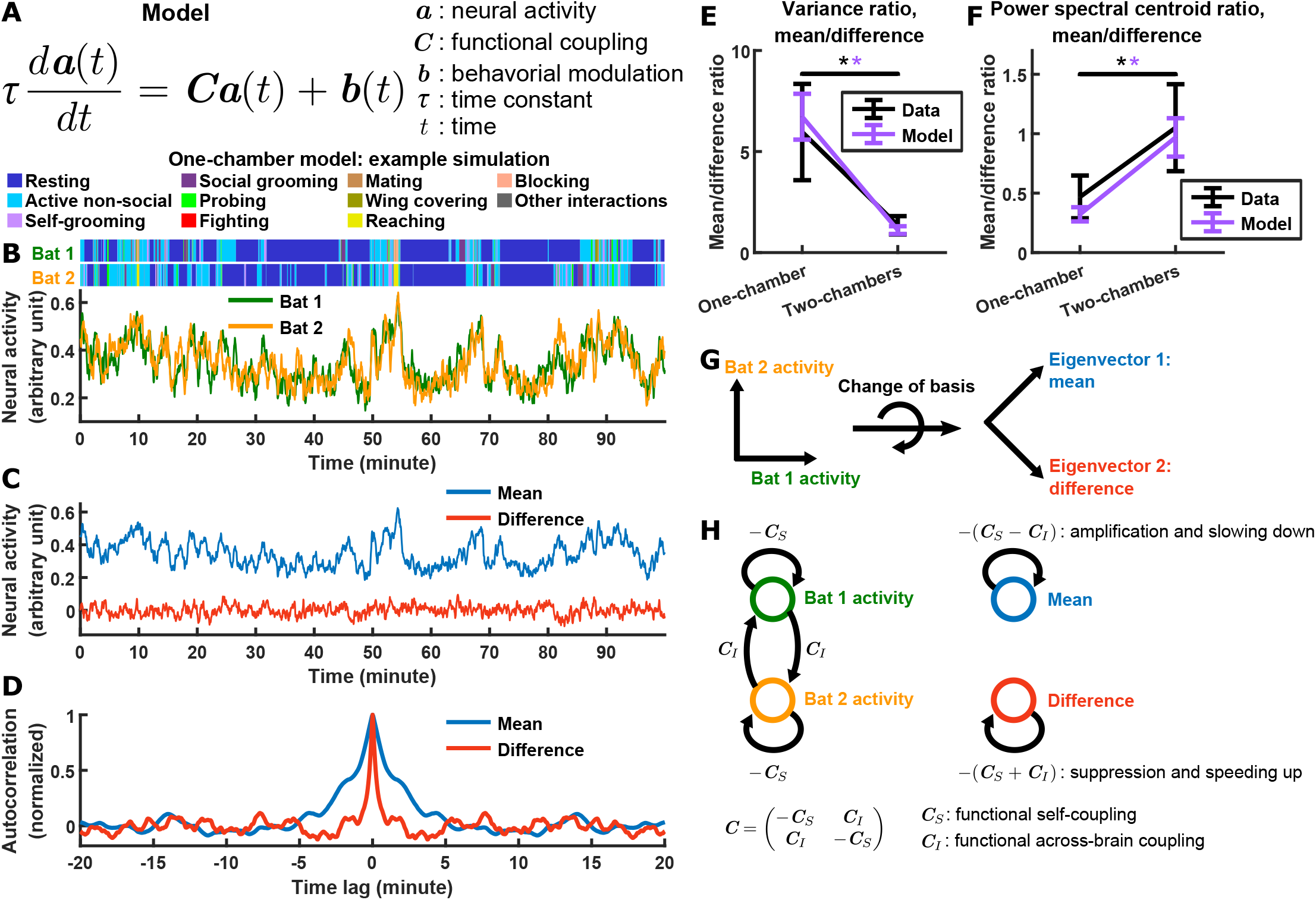
Model explains relationship between difference and mean. (A) The evolving neural activity of two bats are modeled by a linear differential equation. 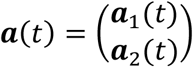 is the activity of bat 1 and bat 2 at time *t*. 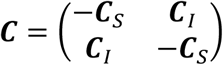 is the functional coupling matrix, where ***C***_*S*_ is the strength of functional self-coupling and ***C***_*I*_ is the strength of functional across-brain coupling (note that functional across-brain coupling obviously should not be interpreted as direct coupling via actual neural connections). 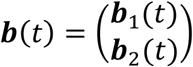 is the modulation of each bat’s activity by its behaviors, where the behaviors are simulated using Markov chains. See Supplementary Information section 4.4 for details. (B) Simulated neural activity and behaviors from an example one-chamber simulation. (C) The simulated activity from (B) plotted as the activity of its mean and difference components. Note the smaller magnitude of the difference compared to the mean. (D) The autocorrelations (peak-normalized) of the mean and difference components shown in (C). The autocorrelations were computed after subtracting from each time series its average over time. Note that the difference varied on faster timescales than the mean. (E)-(F) The average variance ratio (E; mean component variance divided by difference component variance) and average power spectral centroid ratio (F; mean component centroid divided by difference component centroid), for model simulations (purple) or mean normalized 30-150 Hz LFP power from the data (black). The averages were taken across simulations for the model, and across sessions for the data. Error bars denote standard deviations. *, p<0.05, Wilcoxon rank sum test. (G)-(H) Model mechanism. See Supplementary Information section 4.5 for details.Schematic illustrating a change of basis from the activity of each bat to the eigenvectors of the functional coupling matrix ***C***. This corresponds to changing to the basis of mean and difference components. (H) The neural activity variables of the two bats are coupled to each other (left). Changing to the eigenvector basis transforms them into uncoupled variables: the mean and difference components (right). The mean and difference components each provides feedback onto itself, with feedback strengths being the eigenvalues of ***C***, which depends on the functional across-brain coupling ***C***_*I*_. In the one-chamber model, ***C***_*I*_ > 0, so functional across-brain coupling amplifies and slows down the mean component through positive feedback, and suppresses and speeds up the difference component through negative feedback.

This model qualitatively reproduces not only the experimentally observed neural activity patterns (Figure 3B), but also all of the experimentally observed relationships between the magnitudes and timescales of the difference and mean components (Figure 3C-F). Specifically, the difference component had smaller magnitudes and faster timescales than the mean component in the one-chamber model, but not in the two-chambers model (Figure 3C-F).

What mechanisms in the model are responsible for these results? We now analyze the model to answer this (see Supplementary Information section 4.5 for details). The model describes the evolving neural activity of two bats using separate variables to represent the activity of each bat, as in the traditional framework. This is a basis in which the neural activity variables are coupled to each other in the one-chamber model: the activity of bat 1 influences the activity of bat 2, which in turn feeds back on the activity of bat 1. It is much easier to understand how activity evolves if we change to a basis in which the activity variables become uncoupled. This basis consists of the eigenvectors of the functional coupling matrix ***C***. Under the eigenvector basis, the neural activity variables become the mean activity across bats, and the difference in activity between bats (Figure 3G). Thus, the uncoupled activity variables are precisely our variables of interest: the inter-brain mean and difference components. Each of the two components provides feedback onto itself, with feedback strengths being the eigenvalues of ***C***: −(***C***_*S*_ − ***C***_*I*_) for the mean component and −(***C***_*S*_ + ***C***_*I*_) for the difference component. In the two-chambers model, ***C***_*I*_ = 0, so the two components receive equal feedback. In the one-chamber model, ***C***_*I*_ > 0, so functional across-brain coupling acts as positive feedback for the mean component, which amplifies the mean component while slowing it down. On the other hand, functional across-brain coupling acts as negative feedback for the difference component, which suppresses the difference component while speeding it up (Figure 3H). Thus, in the model, a single mechanism—opposite feedback to the inter-brain difference and mean components—contributes to all of our observations relating the magnitudes and timescales of the mean and difference components.

Next, we explored the extent to which alternative models could reproduce the data. In our main model, in addition to functional coupling, coordinated behavioral modulation also contributes to the relationships between mean and difference (Supplementary Information section 4.5; Figure S4). Thus, we studied an alternative model with coordinated behavioral modulation, but without functional across-brain coupling (Supplementary Information section 4.5). We found that such a model cannot reproduce a key aspect of the experimental observations—inter-brain correlation during time periods when the bats engaged in uncoordinated behaviors (Figure S5A-G).

We also explored an alternative mechanism for functional across-brain coupling, using the Kuramoto model (Supplementary Information section 4.6; Strogatz, 2000). The Kuramoto model has been widely used to model diverse synchronization phenomena in physics, chemistry, and biology, including neuroscience (e.g., Breakspear et al., 2010; Cumin and Unsworth, 2007; Schmidt et al., 2015; see also review by Acebrón et al., 2005), and thus we adapted it here to model inter-brain synchronization. In this model, the fluctuating neural activity of the interacting bats are abstracted as oscillators whose phases are dynamically coupled depending on their phase difference. This phase-coupling mechanism is able to reproduce inter-brain correlation and the relative magnitudes of the mean and difference components from the data; however, importantly, it does not reproduce the relative timescales of the mean and difference components from the data (Figure S5H-M).

In summary, our main model, but not the alternative models, provides a parsimonious explanation to the set of robust, but puzzling relationships between inter-brain mean and difference components. Namely, through opposite feedback to the mean and difference components, functional across-brain coupling simultaneously modulates both their magnitude and timescales in opposite directions—the amplification and slowing down of the mean and the suppression and speeding up of the difference are all manifestations of a single mechanism. This insight stemmed from our new framework for examining the inter-brain neural relationship: in the data, the change of basis to the mean and difference components revealed fast inter-brain catch-up riding on top of slow activity covariation across brains; and in the model, the change of basis led to an explanation for these observations. Combined, the data and model suggest a unifying computational mechanism shaping the inter-brain relationship.

### Testing predictions emerging from the model

In addition to explaining existing experimental observations, the model also makes novel predictions about previously unexamined aspects of the inter-brain relationship. We now describe these predictions, and then turn back to the data to test them.

In the model, the neural activity of each bat is a variable. These two activity variables can be thought of as evolving within a two-dimensional space, i.e., a space whose axes are the neural activity of each of the two bats (Figure 3G left). The evolution of activity in this space can be equivalently described using alternative sets of activity variables: for example, by rotating the axes by 45°, the activity variables become the difference and mean components (Figure 3G right). Rotating the axes in this way changes not only the activity variables, but also the strength of functional coupling between the variables. In the one-chamber model, if we rotate the axes smoothly, the strength of functional coupling also changes smoothly (Figure 4A). In particular, the coupling strength is positive and at its maximum when the axes correspond to the activity of each bat, and it decreases to zero as the axes rotate by 45° (corresponding to the mean and difference components), and finally becoming negative as the axes rotate further (Figure 4A). After any amount of rotation, we can calculate the correlation between the activity variables. The model predicts that, as the axes rotate, the correlation between the neural activity variables would mirror the coupling strength (Figure 4B). This effect is not due to the changing behavioral modulation upon axes rotation, as can be seen after regressing out the influence of behavior from the activity (brown curve in Figure 4B). As we turn back to the data, we found that it clearly confirmed the model predictions (Figure 4C). Similarly, data from the two-chambers sessions (Figure 4F) also confirmed the predictions of the two-chambers model (Figure 4D-E), namely, near zero correlation across all axes rotation angles.

**Figure 4.**
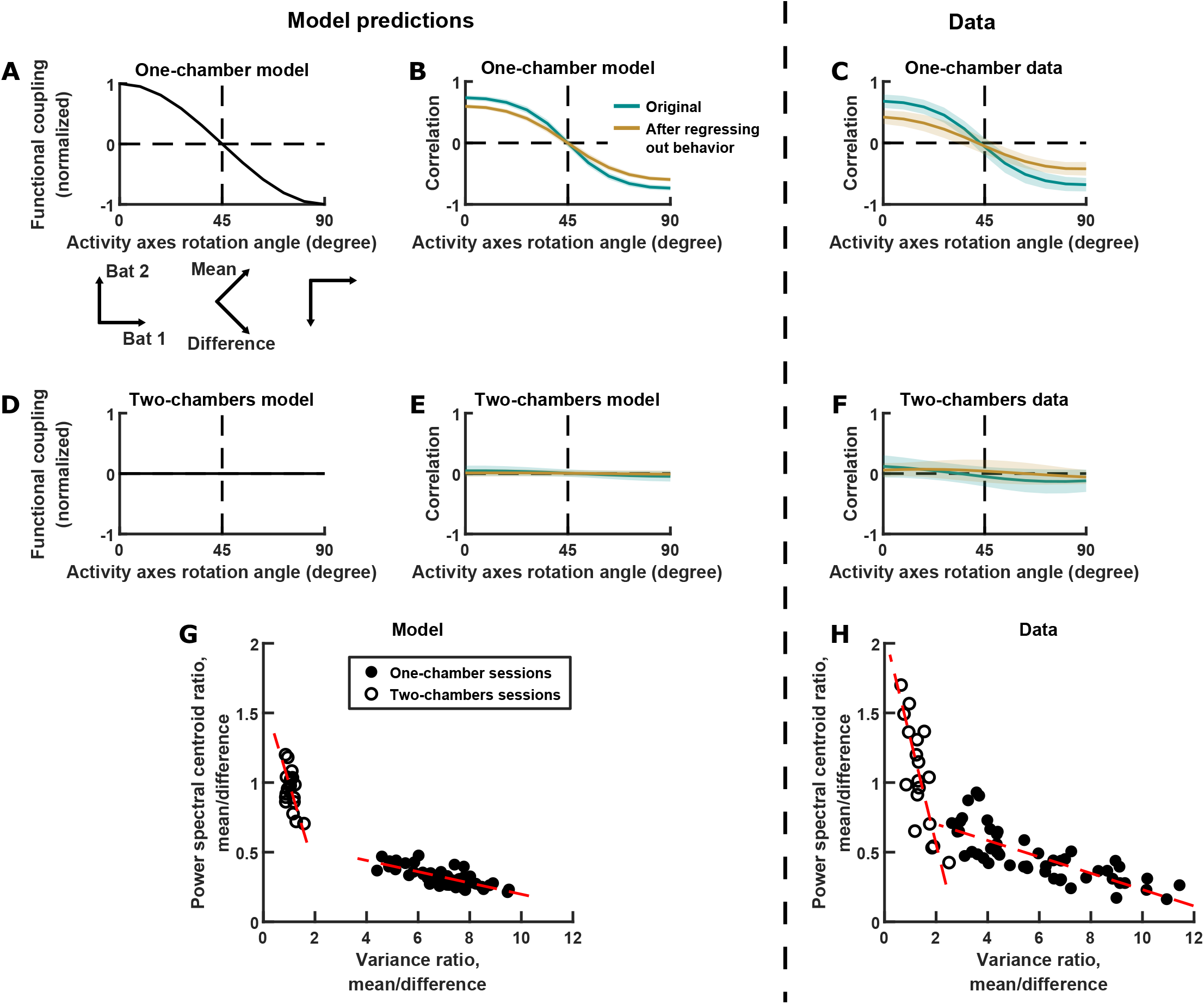
Testing model predictions. (A), (B), (D), (E), and (G) show model results and predictions, and (C), (F), and (H) show the corresponding data. (A) Functional coupling between the activity variables in the one-chamber model, plotted against the orientation of the axes of the activity space that defines the activity variables. When the activity variables correspond to the activity of each bat (0° rotation), functional coupling is at its positive maximum. Functional coupling decreases as the axes rotate, becoming zero when the activity variables correspond to the mean and difference components (45° rotation). Three orientations of the axes are illustrated below the plot. The plotted functional coupling was normalized by the functional coupling at 0° rotation. (B) Correlation between the activity variables in the one-chamber model, plotted against the orientation of the axes of the activity space that defines the activity variables. Correlations were shown both before (teal) and after (brown) regressing out the behaviors of both bats from each activity variable. Shading indicates standard deviation across simulations. Note that the shape of the correlation curves mirrors that of functional coupling shown in (A), and that regressing out behaviors changes the magnitude, but not the shape of the correlation curve. (C) Same as (B), but for data from one-chamber sessions. Shading indicates standard deviation across sessions. Note that the data confirms the model predictions from (B). (D)-(F) Same as (A)-(C), but for the two-chambers model and data. Note that the data again confirms model predictions. (G) Scatter plot of power spectral centroid ratio (mean component centroid divided by difference component centroid) vs. variance ratio (mean component variance divided by difference component variance). Each circle is a simulation (filled: one-chamber model; open: two-chambers model). Red dashed lines are total least squares regression lines. Note that the model predicts linear relationships between the variance ratio and centroid ratio. See Figure S6 and Supplementary Information section 4.8 for detailed analysis of these relationships. (H) Same as (G), but for the data. Note that the prediction from (G) is confirmed by the data.

In the previous sections, we have examined in detail the relationships between the magnitudes and timescales of the inter-brain activity components in single sessions, but the model also makes predictions regarding their relationship across sessions. As described above and in Supplementary Information (section 4.5), functional across-brain coupling acts as positive feedback for the mean component and negative feedback for the difference component, modulating both their magnitudes and timescales in opposite directions (Figure 3H). Thus, the model predicts that the relative magnitudes and relative timescales of the two components are tied together by this mechanism and do not vary independently. This can be seen by plotting the relative magnitudes against the relative timescales for different simulations, which shows linear relationships between them (Figure 4G; see Supplementary Information section 4.8 for detailed analysis of these relationships). Turning to the data, we see similar linear relationships, again confirming the predictions of the model (Figure 4H). It is important to note that these are very specific predictions that were not at all obvious or expected from our previous knowledge of the inter-brain relationship. The fact that they were verified by the data provides strong support for the validity of the model.

In the model, the neural activity of each bat is directly modulated by its own behavior; moreover, it is indirectly modulated by the behavior of the other bat, through functional across-brain coupling. This suggests that the neural activity of each bat should represent the behavior of the other bat independently from encoding its own behavior. To quantify this in the model, we first regressed out the behavior of each bat from its own neural activity, then examined to what extent the residual neural activity encodes the behavior of the other bat, using the receiver operating characteristic (ROC) curve. Importantly, by regressing out each bat’s own behavior, this approach eliminates potential spurious correlation between one bat’s activity and another bat’s behavior caused by any coordinated behaviors between the bats. Using this method, Figure 5A-C shows that the activity of each bat in the model is modulated by the behaviors of the other bat independently of its own behavior. Applying the same approach to the data, we found that the model predictions are again confirmed (Figure 5D-F).

**Figure 5.**
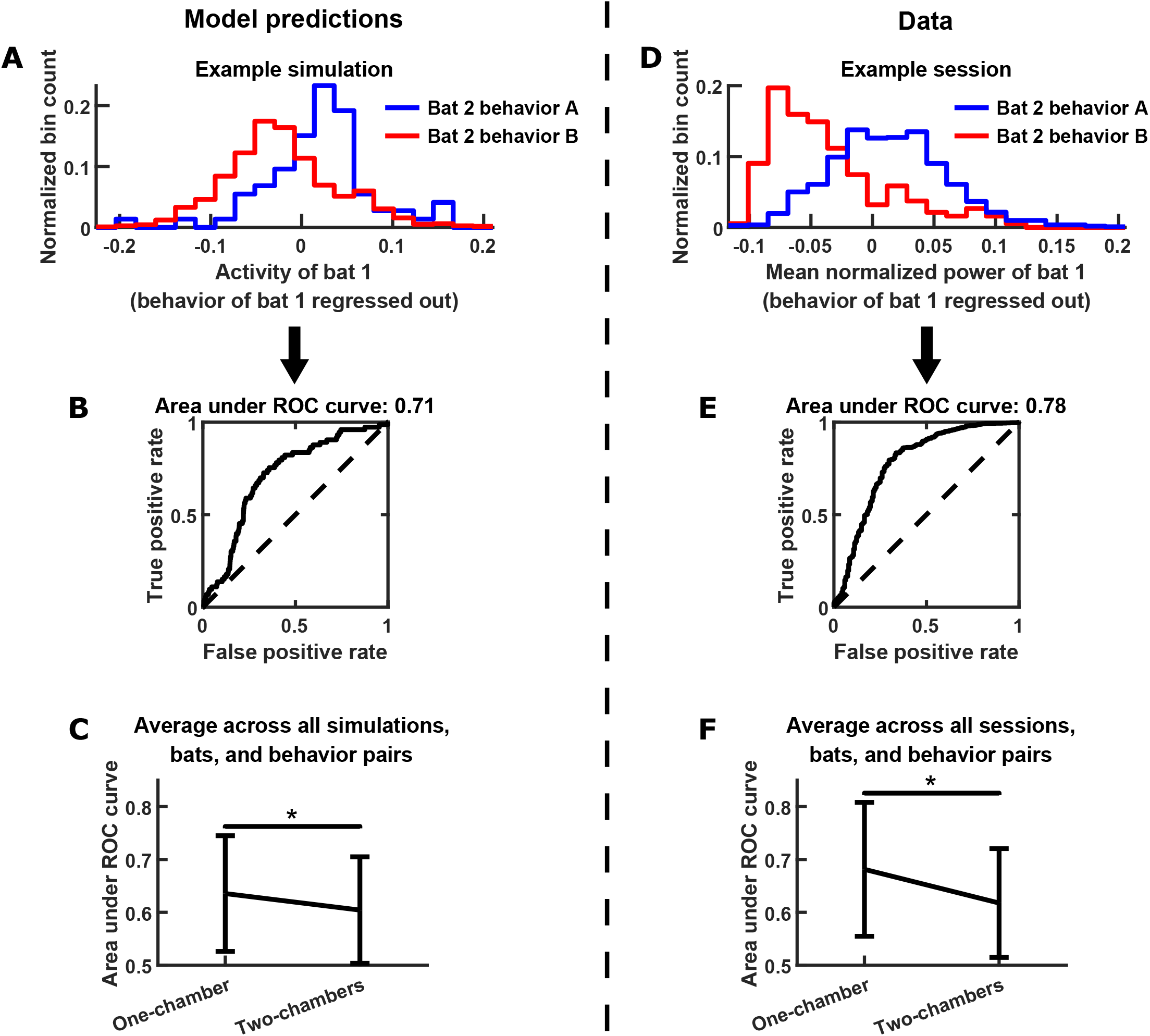
Testing further model predictions. (A)-(C) show model results and predictions, and (D)-(F) show the corresponding data. (A) Distributions of the neural activity of bat 1 conditioned on the behavior of bat 2. Bat 1’s behavior has been regressed out of its neural activity. Two distributions are shown from an example simulation, for two example behaviors by bat 2 (behavior A: probing; behavior B: resting). Note that bat 1’s activity encodes bat 2’s behavior independently of its own behavior. (B) The extent to which bat 1’s activity encodes the two behaviors of bat 2 in the example from (A) is quantified using an ROC curve, which illustrates the discriminability of the two distributions from (A). The area under the ROC curve is indicated above the plot (larger area indicates better discriminability). (C) Area under the ROC curve is averaged across all simulations, bats, and behavior pairs, separately for one-chamber simulations and two-chambers simulations. When using one bat’s activity to discriminate the other bat’s behavior, discriminability is significantly higher in one-chamber simulations. Error bars denote standard deviations. *, p<0.05, Wilcoxon rank sum test. (D)-(F) Same as (A)-(C), but for mean normalized 30-150 Hz LFP power from the data. For the example in (D), behaviors A and B are active non-social and social grooming, respectively. Note that the data is consistent with model predictions.

In summary, multiple predictions of the model found support in the data, underlining its explanatory power.

### Modeling inter-brain neural relationship during group social interactions

Our experiments and model were exclusively focused on social interactions between two individuals. However, social interactions also occur among larger groups in many species, including the group-living Egyptian fruit bats (Herzig-Straschil and Robinson, 1978; Kwiecinski and Griffiths, 1999). While the inter-brain relationship between two interacting individuals have been studied extensively, the inter-brain relationship among larger social groups have received far less attention (but see Dikker et al., 2017). Therefore, we extended our model of two interacting bats to larger groups, in order to offer direct predictions on the group inter-brain relationship, which can be directly tested using recently established approaches (Kingsbury et al., 2019; Zhang and Yartsev, 2019).

To extend the two-bat model to interactions among a group of *n* bats, we used the same equation shown in Figure 3A, except that the activity (***a***) and behavioral modulation (***b***) are now *n*-dimensional rather than 2-dimensional, and the functional coupling matrix ***C*** is similarly *n* × *n*. As in the two-bat model, the activity of each bat in the *n*-bat model is influenced by the same functional self-coupling −***C***_*S*_, and the activity of each pair of bats are functionally coupled with the same positive coupling ***C***_*I*_.

The mechanism that governs the two-bat model naturally generalizes to the *n*-bat model (see Supplementary Information section 4.10 for details). For the *n*-bat model, the mean component corresponds to the mean activity across all bats. On the other hand, a difference subspace takes the place of the difference component of the two-bat model. In *n*-bat activity space, the difference subspace is the (*n* − 1)-dimensional subspace orthogonal to the direction of the mean component (Figure 6B). This subspace contains all inter-brain activity patterns that correspond to activity differences across any of the brains. Similar to the two-bat model, functional across-brain coupling acts as positive and negative feedback to the *n*-bat mean component and the difference subspace, respectively, amplifying and slowing down the *n*-bat mean component while suppressing and speeding up activity patterns in the difference subspace. This gives rise to predictions that can be tested experimentally. Specifically, the model predicts that activity patterns in the difference subspace would have smaller magnitudes and faster timescales than the *n*-bat mean component (Figure 6C-D). Moreover, similar to the two-bat model (see Figure 4A-B), correlation between activity variables depends on their functional coupling. In particular, the positive functional across-brain coupling between the activity of different bats would give rise to positive inter-brain correlations; on the other hand, the mean component is not functionally coupled with activity patterns in the difference subspace, so they would therefore be uncorrelated (Figure 6E).

**Figure 6.**
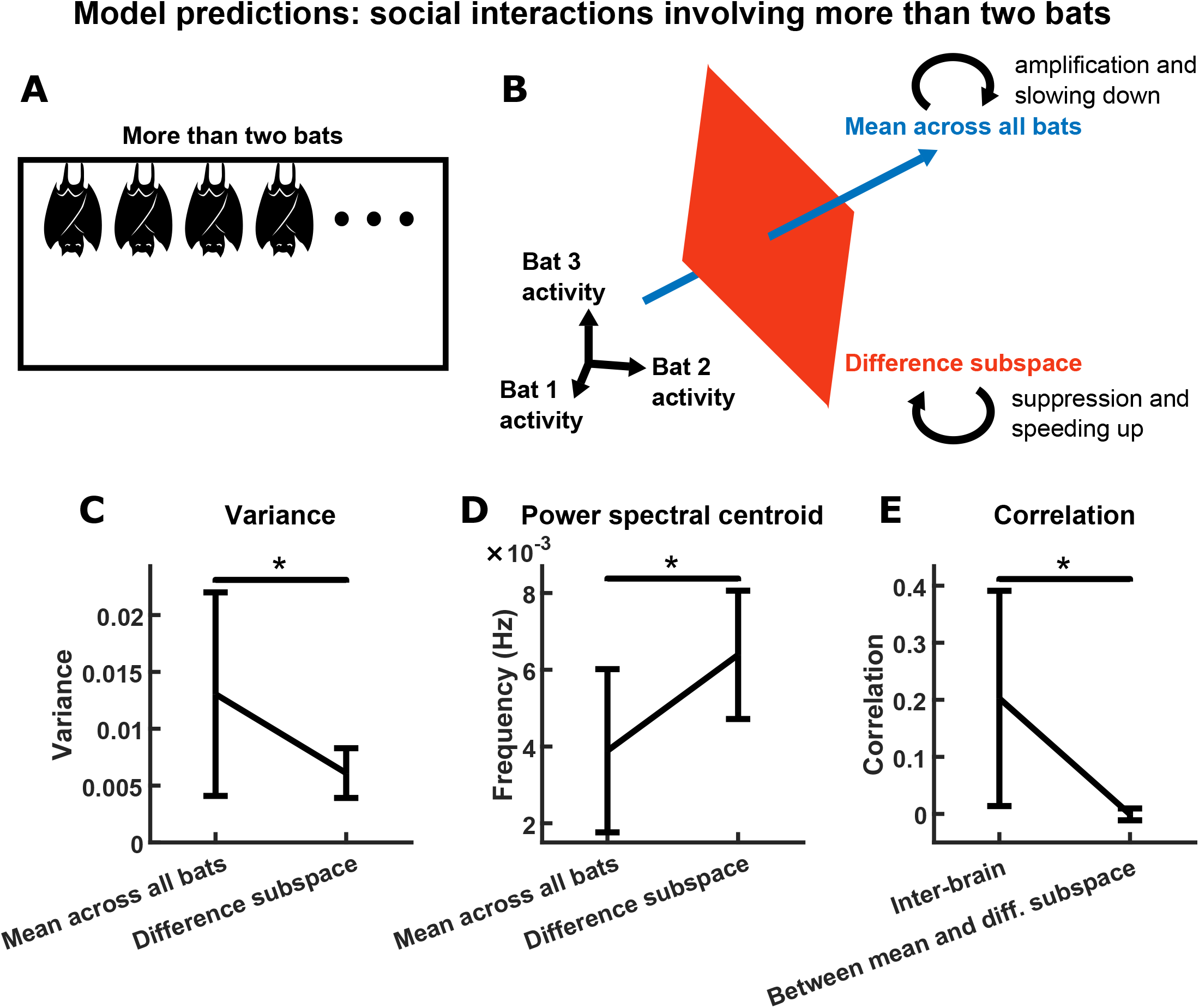
Model predictions for group inter-brain relationship. (A) The two-bat model can be generalized to model the neural activity of a group of more than two socially interacting bats (the *n*-bat model). (B) The mechanism underlying the *n*-bat model, illustrated for the case of *n* = 3 bats (three was chosen only because higher-dimensional spaces cannot be illustrated). Shown are the mean component and the difference subspace in the activity space of *n* bats. The mean component, corresponding to the mean activity across all bats, is a vector pointing towards the first octant. The difference subspace is the (*n* − 1)-dimensional subspace orthogonal to the direction of the mean component, and it contains all inter-brain activity patterns that correspond to activity differences across brains. Similar to the two-bat model, functional across-brain coupling acts as positive feedback to the *n*-bat mean component, amplifying it and slowing it down; and as negative feedback to activity patterns in the difference subspace, suppressing them and speeding them up. (C)-(E) Model predictions from simulations of an *n*-bat model where *n* = 4. (C) Model predictions on the variance of the mean component and the average variance of the difference subspace (total variance in the difference subspace divided by *n* − 1). Plotted are averages (± standard deviation) across simulations. (D) Model predictions on the power spectral centroid of the mean component and the average power spectral centroid of the difference subspace (averaged across the power spectral centroids of activity in 1000 random directions in the difference subspace for a given simulation). Plotted are averages (± standard deviation) across simulations. (E) The model predicts positive inter-brain correlation and zero average correlation between the mean component and activity patterns in the difference subspace. Here, for each simulation, inter-brain correlation was averaged across all pairs of bats, and average correlation was calculated between the mean component and activity in 1000 random directions in the difference subspace. Plotted are averages (± standard deviation) across simulations. *, p<0.05, Wilcoxon signed rank test.

Thus, extension of our two-bat model to group interactions generated explicit predictions regarding the group inter-brain relationship, specifically on the magnitudes, timescales, and correlations of inter-brain activity components (Figure 6C-E). These predictions remain to be tested experimentally, which is now possible with the recent development of techniques to record simultaneously from multiple interacting bats (Zhang and Yartsev, 2019). Testing these predictions will be an important step towards understanding group inter-brain relationships.

## Discussion

Influential theories have posited that functional across-brain coupling is a key determinant of neural activity underlying social interactions (Hasson et al., 2012; Hasson and Frith, 2016). Our results support these theories, and suggest a specific computational mechanism through which functional coupling acts during social interactions—opposite feedback to difference and mean components—which shapes the inter-brain neural relationship. Furthermore, the mechanism we uncovered has broad explanatory power, and can account for a range of previously observed features of the inter-brain neural relationship, as we discuss below.

First, it is well known that neural activity is correlated between the brains of socially interacting individuals, in mice, bats, and humans (e.g., Dikker et al., 2014; Kingsbury et al., 2019; Kinreich et al., 2017; Levy et al., 2017; Montague et al., 2002; Piazza et al., 2020; Spiegelhalder et al., 2014; Stolk et al., 2014; Zadbood et al., 2017; Zhang and Yartsev, 2019). Through negative feedback to the difference component, functional across-brain coupling gives rise to an inter-brain catch-up phenomenon that keeps the inter-brain difference small. At the same time, positive feedback to the mean component amplifies the common activity covariation between brains, so that the neural activity in each is dominated by their common activity pattern. Together, these would result in the inter-brain correlation observed during social interactions in diverse species.

Second, in a shared social environment, neural correlation across brains is above and beyond what would be expected from behavioral coordination between individuals (Kingsbury et al., 2019; Piazza et al., 2020; Zhang and Yartsev, 2019). This can be understood because, through opposite feedback to the difference and mean components, functional across-brain coupling contributes to correlation independently from behavioral coordination. As a consequence of this, even in the absence of behavioral coordination, the opposite feedback results in correlation across brains in the model (Figure S5A-G), consistent with previous experimental observations (Kingsbury et al., 2019; Piazza et al., 2020; Zhang and Yartsev, 2019).

Third, the neural correlation between socially interacting individuals varies as a function of timescale: correlation across brains is higher for activity at slower timescales, and lower for activity at faster timescales (Kingsbury et al., 2019; Zhang and Yartsev, 2019). This would be a natural consequence of the opposite feedback mechanism. Positive feedback to the mean amplifies it and slows it down, while negative feedback to the difference suppresses it and speeds it up. Thus, at slower timescales, the activity of the two brains is dominated by the mean component, i.e., the activity of the two brains are very similar at slower timescales and thus have higher correlations. On the other hand, at faster timescales, the increasing presence of the difference component lowers the correlation across brains.

In summary, our model offers a particularly parsimonious explanation for diverse aspects of the inter-brain neural relationship, including: (1) neural correlation across brains; (2) neural correlation beyond behavioral coordination; (3) neural correlation as a function of timescale; (4) the relative magnitudes and (5) relative timescales of inter-brain mean and difference components (Figure 3E-F); (6) correlation between activity variables under rotated activity bases (Figure 4B-C); (7) the relationship between the magnitudes and timescales of activity components across sessions (Figure 4G-H); and (8) modulation of one bat’s neural activity by the behavior of another bat (Figure 5). Importantly, there is no reason a priori to suspect that these disparate observations are all related; only in light of the model is it apparent that they are all manifestations of a single underlying computational mechanism. Collectively, these results thus suggest opposite feedback to the difference and mean as a unifying computational mechanism governing the inter-brain relationship during social interactions. This points to a major avenue of future research: elucidating the biological implementation of this computational mechanism. Such research will likely involve studying the distributed neural circuits mediating sensorimotor interactions and attentional processes during social interactions, to understand how they might functionally act as opposite feedback to the inter-brain difference and mean components.

## Acknowledgments

We thank E. S. Sevilla, A. Raha, J. Chau, K. Moi, N. Juthani, M. Zuercher, L. Kasraie, and C. Tran for behavioral annotation; S. A. Afjei for histology; L. Jiang for machining; B. Olshausen, W. Liberti, M. Rose and members of the Yartsev Lab for discussion and comments on the manuscript; C. Ferrecchia and G. Lawson for veterinary oversight; the staff of the Office of Laboratory Animal Care for support with animal husbandry and care. This research was supported by NIH (DP2-DC016163), the New York Stem Cell Foundation (NYSCF-R-NI40), the Alfred P. Sloan Foundation (FG-2017-9646), the Brain Research Foundation (BRFSG-2017-09), National Science Foundation (NSF-1550818), the Packard Fellowship (2017-66825), the Klingenstein-Simons Fellowship, the Pew Charitable Trust (00029645), and the Dana Foundation (to M.M.Y.).

## Author Contributions

W.Z. collected the data and performed the analysis and modeling, with help and feedback from M.M.Y. M.M.Y. supervised the entire study. W.Z. and M.M.Y. wrote the manuscript.

## Supplementary Figure Legends

**Figure S1.**
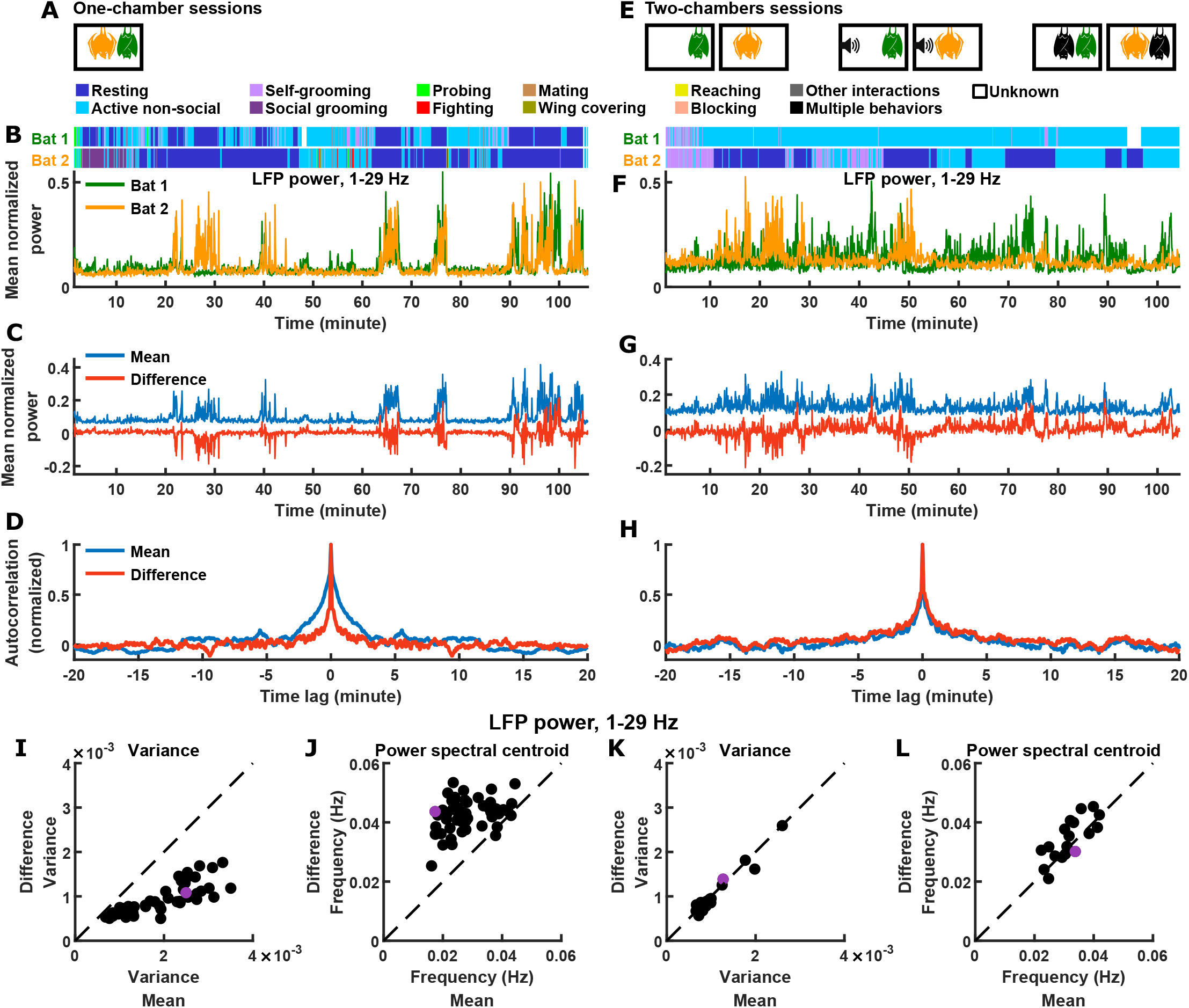
Inter-brain difference and mean components: 1-29 Hz LFP power. Same figure format as Figure 1A-L. (A) One-chamber sessions: simultaneous neural recording from pairs of bats engaged in natural social interactions. (B) Mean normalized LFP power in the 1-29 Hz band (Supplementary Information sections 3.7, 3.9), averaged across all channels for each bat, on an example one-chamber session. Shown above are the annotated behaviors. (C) The mean and difference components of the activity of the two bats shown in (B). (D) Autocorrelations (peak-normalized) of the mean and difference activity components shown in (C). The autocorrelations were computed after subtracting from each time series its average over time. Note that the difference component varied on faster timescales than the mean component. (E) Two-chambers sessions: simultaneous neural recording from the same bats from the one-chamber sessions freely behaving in separate, identical chambers. (F)-(H) Same as (B)-(D), but for an example two-chambers session. (I)-(J) Variance (I) and power spectral centroid (J) of mean normalized 1-29 Hz LFP power, for the mean and difference components. Each dot is one one-chamber session (the purple dot is the session shown in (B)-(D)). Variance quantifies activity magnitude, and power spectral centroid quantifies timescale (higher centroids mean faster timescales). Note that, on one-chamber sessions, the difference component tends to be smaller and faster than the mean component. The dotted lines are unity. Note that the power spectral centroid was calculated from time series of mean normalized LFP power (e.g. as plotted in (C)), not from time series of LFP itself. (K)-(L) Same as (I)-(J), but for two-chambers sessions. The purple dot is the session shown in (F)-(H). The mean and difference components have comparable magnitudes and timescales on the two-chambers sessions.

**Figure S2.**
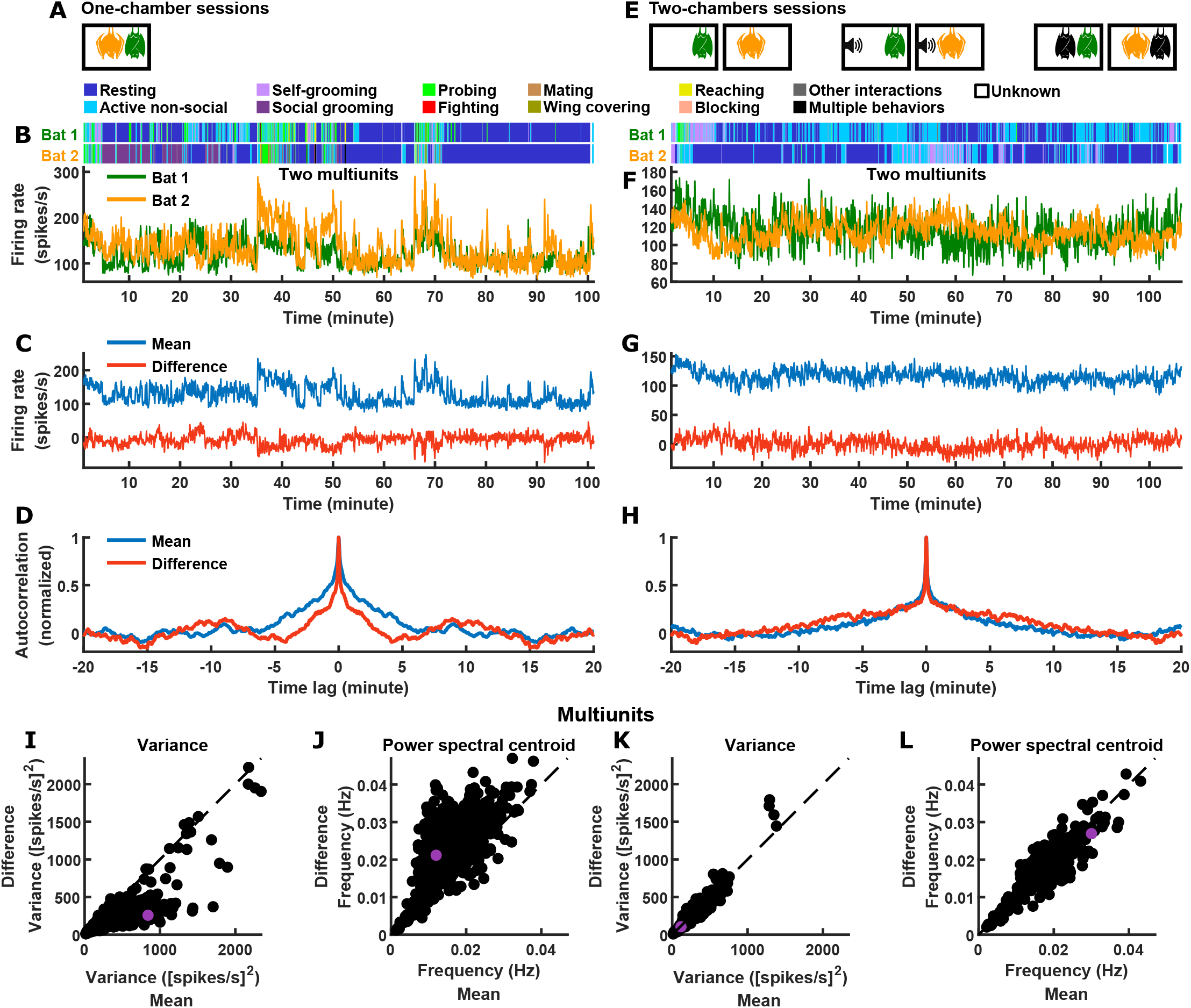
Inter-brain difference and mean components: multiunit activity. Same figure format as Figure 1A-L. (A) One-chamber sessions: simultaneous neural recording from pairs of bats engaged in natural social interactions. (B) Activity of two multiunits, one from each brain, on an example one-chamber session. Shown above are the annotated behaviors. (C) The mean and difference components of the activity of the two bats shown in (B). (D) Autocorrelations (peak-normalized) of the mean and difference activity components shown in (C). The autocorrelations were computed after subtracting from each time series its average over time. Note that the difference component varied on faster timescales than the mean component. (E) Two-chambers sessions: simultaneous neural recording from the same bats from the one-chamber sessions freely behaving in separate, identical chambers. (F)-(H) Same as (B)-(D), but for an example two-chambers session. (I)-(J) Variance (I) and power spectral centroid (J) of multiunit activity, for the mean and difference components. Each dot is one pair of multiunits (one from each brain) from one-chamber sessions (the purple dot is the pair shown in (B)-(D)). Variance quantifies activity magnitude, and power spectral centroid quantifies timescale (higher centroids mean faster timescales). Note that, on one-chamber sessions, the difference component tends to be smaller and faster than the mean component. The dotted lines are unity. (K)-(L) Same as (I)-(J), but for two-chambers sessions. The purple dot is the pair of multiunits shown in (F)-(H). The mean and difference components have comparable magnitudes and timescales on the two-chambers sessions.

**Figure S3.**
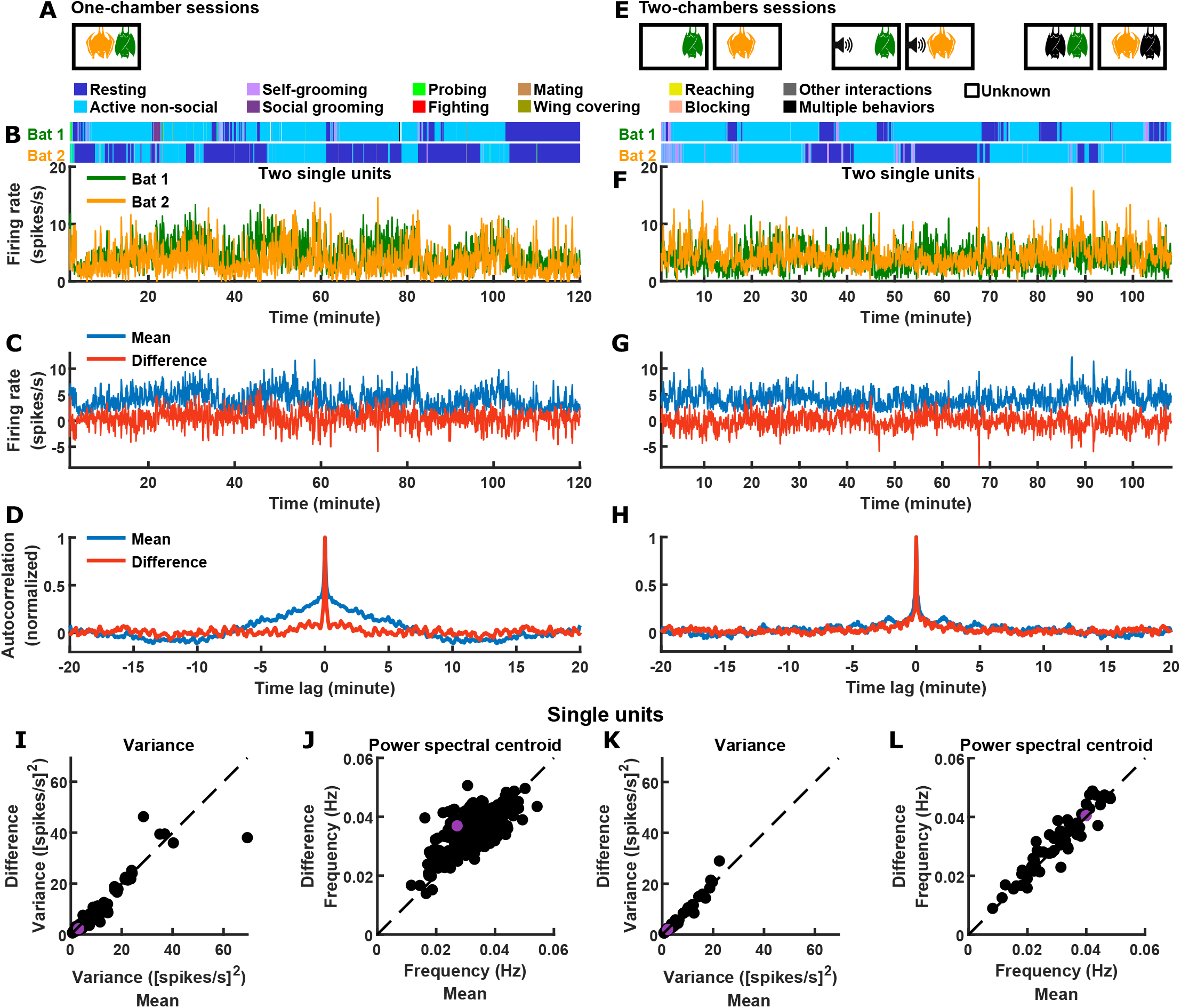
Inter-brain difference and mean components: single unit activity. Same figure format as Figure 1A-L. (A) One-chamber sessions: simultaneous neural recording from pairs of bats engaged in natural social interactions. (B) Activity of two single units, one from each brain, on an example one-chamber session.Shown above are the annotated behaviors. (C) The mean and difference components of the activity of the two bats shown in (B). (D) Autocorrelations (peak-normalized) of the mean and difference activity components shown in (C). The autocorrelations were computed after subtracting from each time series its average over time. (E) Two-chambers sessions: simultaneous neural recording from the same bats from the one-chamber sessions freely behaving in separate, identical chambers. (F)-(H) Same as (B)-(D), but for an example two-chambers session. (I)-(J) Variance (I) and power spectral centroid (J) of single unit activity, for the mean and difference components. Each dot is one pair of single units (one from each brain) from one-chamber sessions (the purple dot is the pair shown in (B)-(D)). Variance quantifies activity magnitude, and power spectral centroid quantifies timescale (higher centroids mean faster timescales). The dotted lines are unity. (K)-(L) Same as (I)-(J), but for two-chambers sessions. The purple dot is the pair of single units shown in (F)-(H).

**Figure S4.**
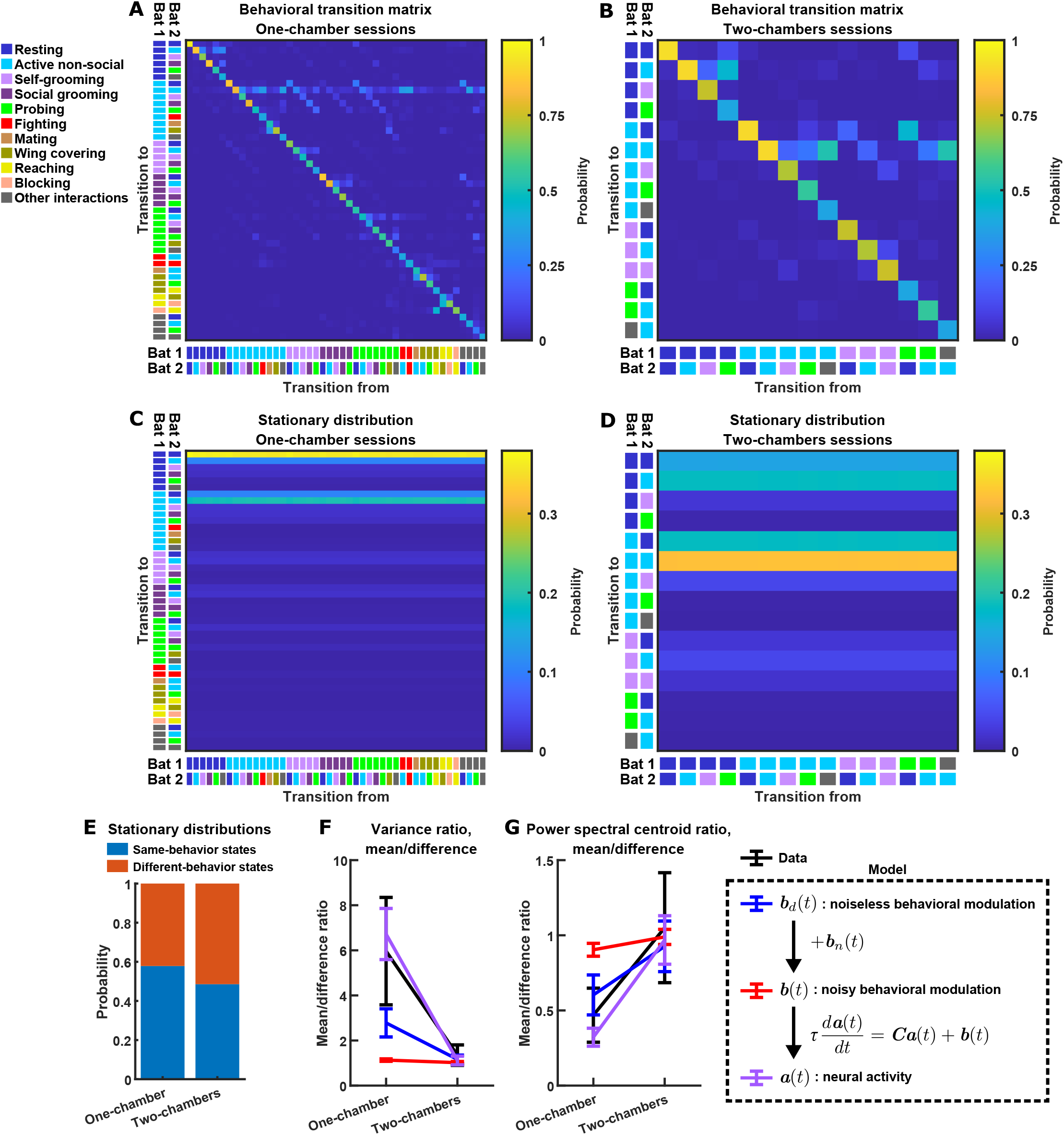
Modeling behavior and behavioral modulation of difference and mean components. (A)-(B) Empirical behavioral transition frequency matrix for the one-chamber (A) and two-chambers (B) sessions, used to simulate behaviors using Markov chains (Supplementary Information section 4.4). A behavioral state consists of the behaviors of both bats: e.g. one state could be “bat 1 resting, bat 2 self-grooming.” Each matrix shows the frequencies of transition from one state at a given time to another state after a passage of 2.5 s (the time step size of the Markov chains): the color of the matrix element at row *i* and column *j* denotes the frequency of transitions from state *j* to state *i*. The states are labeled by color to the left and bottom of the matrices. Here the diagonal elements (self-transitions) are larger than the off-diagonal elements (transitions between states) because of the small time step size of the Markov chain. The transition frequencies were calculated after pooling the behavioral data from all one-chamber (A) or two-chambers (B) sessions. (C) Multi-step transition matrix showing the transition probability from one state at a given time to another state after a passage of 5 minutes, in the one-chamber model. This matrix was obtained by raising the single-step transition matrix in (A) to the 120th power. Note that the columns of the multi-step transition matrix are approximately identical, showing that the Markov chain approaches the stationary distribution represented by the columns within ∼5 minutes, i.e. the distribution of states no longer changes with time ∼5 minutes into a simulated session. (D) Same as (C) but for the two-chambers model. (E) The probability of same-behavior states and different-behavior states in the stationary distributions of the one-chamber and two-chambers models. A same-behavior state is defined as a state where the two bats engage in the same behavior, and a different-behavior state is one where the two bats engage in different behaviors. In the stationary distribution of the two-chambers model, same-behavior states (probability: 0.49) are about equally likely as different-behavior states. In the stationary distribution of the one-chamber model, same-behavior states (probability: 0.58) are somewhat more likely than different-behavior states. (F)-(G) The average variance ratio (F; mean component variance divided by difference component variance) and average power spectral centroid ratio (G; mean component centroid divided by difference component centroid), for variables from the model (blue, red, and purple) and for mean normalized 30-150 Hz LFP power from the data (black). The model variables (Supplementary Information sections 4.4-4.5) include simulated noiseless behavioral modulation (blue), simulated noisy behavioral modulation (red), and simulated neural activity (purple). Note that in the one-chamber model, the noiseless behavioral modulation of the mean component is larger and slower than that of the difference component (blue). Behavioral modulation noise decreases the differential modulation of the mean and difference components (red). However, even when the noise strongly reduces differential behavioral modulation of the mean and difference components, the experimentally observed levels of relative activity magnitudes and timescales can still result (purple), due to functional across-brain coupling (Figure 3G-H and Supplementary Information section 4.5). All plotted averages were taken across simulations for the model, or across sessions for the data. The data (black) and simulated activity (purple) are replotted from Figure 3E-F. Error bars denote standard deviations. See Supplementary Information section 4.5 for details.

**Figure S5.**
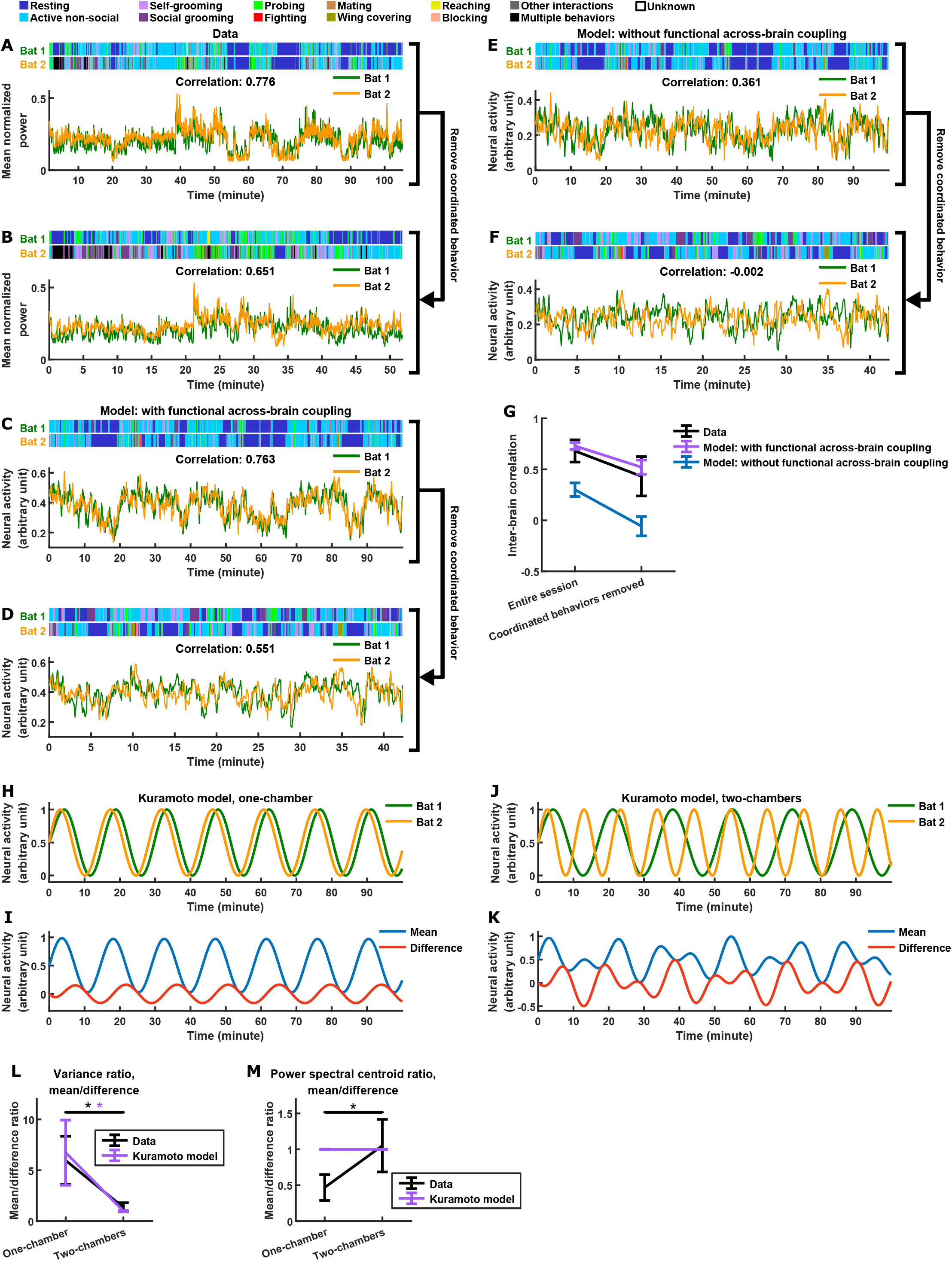
Alternative models. (A)-(G) Comparing models with and without functional across-brain coupling. See Supplementary Information section 4.5 for details. (A) Mean normalized LFP power in the 30-150 Hz band, averaged across all channels for each bat, on an example one-chamber session. Shown above are the annotated behaviors. Note that the activity of the two bats are highly correlated (correlation coefficient indicated above the activity traces). (B) The same data as in (A), after removing all time periods of coordinated behavior (i.e. time periods when both bats engaged in the same behavior). Inter-brain correlation was re-calculated and indicated above the activity traces. Note that the inter-brain correlation is lower than in (A), but still highly positive. (C)-(D) Same as (A)-(B), but for simulated neural activity and behaviors from an example one-chamber simulation of our main model. Note that inter-brain correlation remained after removing time periods of coordinated behavior, reproducing the experimental result. (E)-(F) Same as (C)-(D), but simulated without functional across-brain coupling. Specifically, the same simulated behavior and behavioral modulation ***b***(*t*) (including the same instantiation of behavioral modulation noise ***b***_*n*_(*t*)) was used as in (C)-(D); the difference from (C)-(D) is that the strength of functional across-brain coupling ***C***_*I*_ was set to 0. Note that inter-brain correlation disappeared after removing time periods of coordinated behavior, unlike the experimental result. (G) Inter-brain correlation over entire sessions or after removing coordinated behavior, for data (black; mean normalized LFP power in the 30-150 Hz band), the main model (purple; with functional across-brain coupling), and the model without functional across-brain coupling (blue). The simulations of the two models were done in pairs: each simulation of the model without functional across-brain coupling used the same behavioral modulation (including behavioral modulation noise) as one simulation of the main model, as done in (C)-(F). Plotted are averages (± standard deviation) across experimental sessions or simulations. Note that functional across-brain coupling is required in the model to reproduce persisting inter-brain correlation after the removal of coordinated behaviors. (H)-(M) A model with an alternative form of functional across-brain coupling: the Kuramoto model. In this model, the neural activity of the interacting bats are abstracted as oscillators whose phases are dynamically coupled depending on their phase difference. See Supplementary Information section 4.6 for details. (H)-(I) Example simulation of the one-chamber Kuramoto model, shown as the activity of the two bats (H) and as the mean and difference between bats (I). The model exhibits inter-brain correlation, as well as a larger magnitude for the mean component compared to the difference component. However, the difference and mean components have the same timescale in this model (note that the difference and mean undergo the same number of oscillations per unit time). (J)-(K) Same as (H)-(I), but for the two-chambers Kuramoto model. Here, the difference and mean components have similar magnitudes and timescales. (L)-(M) The average variance ratio (L; mean component variance divided by difference component variance) and average power spectral centroid ratio (M; mean component centroid divided by difference component centroid), for Kuramoto model simulations (purple) or mean normalized 30-150 Hz LFP power from the data (black). The averages were taken across simulations for the model, and across sessions for the data. Note that the Kuramoto model reproduces the relative magnitudes, but not the relative timescales, from the data. The results for data (black) are replotted from Figure 3E-F. Error bars denote standard deviations. *, p<0.05, Wilcoxon rank sum test.

**Figure S6.**
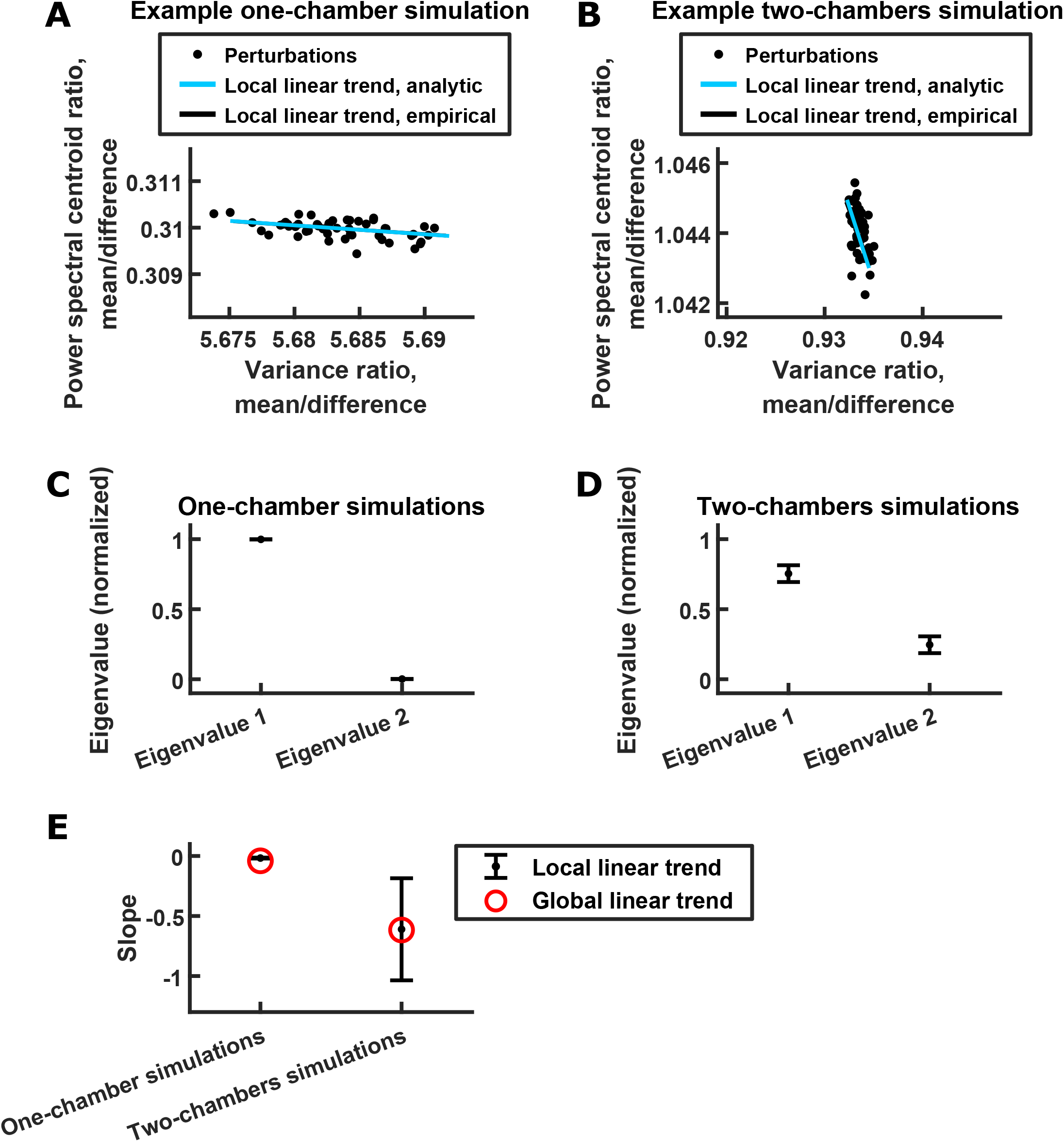
Analysis of linear relationship between power spectral centroid ratio and variance ratio. See Supplementary Information section 4.8 for details. (A) Scatter plot of power spectral centroid ratio (mean component centroid divided by difference component centroid) vs. variance ratio (mean component variance divided by difference component variance), after small random perturbations to the behavioral modulation for a single one-chamber simulation. Each dot is the result of one random perturbation. The blue line is the analytic estimated direction of the linear trend (Supplementary Information section 4.8), and the black line is the empirical direction of the linear trend calculated from 1000 random perturbations (of which a random subset of 50 are plotted, for clarity). The black line is not visible because it is overlapped by the blue line. Note that a clear linear trend is present despite behavioral modulation being randomly perturbed. (B) Same as (A) but for a two-chambers simulation. (C) For a given one-chamber simulation, we estimated the eigenvalues of the covariance matrix of power spectral centroid ratio and variance ratio resulting from random perturbations to the behavioral modulation of that simulation (Supplementary Information section 4.8). Each eigenvalue was then normalized by the sum of the eigenvalues. Plotted are the averages (± standard deviation) of the normalized eigenvalues across simulations, with eigenvalue 1 being the larger eigenvalue from each simulation. Note that one eigenvalue was larger than the other across simulations, indicating a linear trend. (D) Same as (C) but for two-chambers simulations. Note that, again, one eigenvalue tended to be larger than the other. (E) The slopes of the global (red) and local (black) linear relationships between power spectral centroid ratio and variance ratio. The global slopes are the slopes of the linear fit across simulations (red dashed lines in Figure 4G). The local slopes are the slopes of the blue lines in (A)-(B): for each simulation, the local slope was calculated from the leading eigenvector of the covariance matrix of power spectral centroid ratio and variance ratio resulting from random perturbations (Supplementary Information section 4.8). The average local slopes across simulations were then plotted (error bars denote standard deviations). Note that the local slopes, which are not influenced by systematic variations of behavioral modulation across simulations, are consistent with the global slopes.

## Supplementary Information

### 1 CONTACT FOR REAGENT AND RESOURCE SHARING

Further information and requests for resources and reagents should be directed to and will be fulfilled by Michael M. Yartsev.

### 2 EXPERIMENTAL MODEL AND SUBJECT DETAILS

Neural activity was recorded from four adult male Egyptian fruit bats, *Rousettus aegyptiacus* (weight 162-186 g at implantation). 18 bats (11 males, 7 females) were utilized for behavioral experiments only. All bats were wild-caught, and hence their precise age could not be identified (this species of bats is very long-lived, with a maximum reported longevity of 25 years; Kwiecinski and Griffiths, 1999). Before the start of experiments, bats were housed in large communal rooms at the bat breeding colony of UC Berkeley. After the start of experiments, bats were single-housed in cages in a humidity- and temperature-controlled room. The bats were kept on a 12-hour reversed light-dark cycle, and all experiments were conducted during the dark cycle. All experimental procedures were approved by the Animal Care and Use Committee of UC Berkeley.

### 3 METHOD DETAILS

The data analyzed in this study have been previously published in Zhang and Yartsev (2019). Unless otherwise specified, all data processing and analysis was performed using MATLAB (MathWorks).

#### 3.1 Experimental setup and details

All experiments were conducted inside 40.6 × 33.7 × 52.1 cm (length X width X height) cages, which had netting on top that allowed bats to hang. Each cage was placed inside a 64.8 × 61 × 64.8 cm (length X width X height) chamber. Fans circulated air between the inside and outside of the chambers. The experiments were conducted in the dark. The only lights in the chambers were infrared lights to allow video recording. Video was recorded throughout the experimental sessions using one or two high-speed infrared cameras (Flea3, FLIR) at 100 frames/s. Ultrasonic microphones (USG Electret Ultrasound Microphone, Avisoft Bioacoustics; frequency range: 10-120 kHz) were used to record audio throughout the experimental sessions. Infrared light-emitting diodes (LEDs) on the wireless neural recording systems (see below) were flashed at intervals of ∼20 minutes during recording sessions, which were captured by the video cameras and used to synchronize neural and video recordings. Transistor-transistor logic pulses were sent using UltraSoundGate Player 216H (Avisoft Bioacoustics) simultaneously to the wireless neural recording systems of both bats as well as to the audio recording system (UltraSoundGate 416H, Avisoft Bioacoustics), synchronizing neural recording from both bats and the audio recording. All experiments took place in an electromagnetically and acoustically shielded room (IAC Acoustics).

Before the first recording session, each bat used in the experiment was allowed to familiarize itself with the recording environment. This was done in ∼5 familiarization sessions per bat, where a bat freely interacted with another bat for ∼100 minutes per session. One to two of those ∼5 familiarization sessions involved interaction between the same bats that were to be used in the upcoming simultaneous two-bat neural recording experiment.

In total, we recorded 52 one-chamber sessions and 18 two-chambers sessions. In all sessions, bats were allowed to freely behave without intervention or constraint from experimenters. Among the two-chambers sessions, there were 5 sessions of free behavior without playback or interaction partners, 8 sessions of free behavior with playback of bat calls, and 5 sessions of free behavior with non-implanted interaction partners. Identical cages, chambers, and recording setups were used for all one-chamber and two-chambers sessions. Sessions lasted on average 105 ± 6 minutes (mean ± STD). On 31 of the one-chamber sessions, a non-implanted third bat was introduced in the middle of the session. The time of introduction was on average 44.1 ± 14.2 minutes (mean ± STD) from the beginning of the session. The bats that were introduced as the third bat included 11 males and 7 females.

On the two-chambers sessions with playback, bat social calls were played using ultrasound speakers (Vifa, Avisoft Bioacoustics; frequency range: 1-120 kHz). The contents of the playback were identical for the two chambers and were delivered simultaneously to both. On each playback session, a set of different calls were played. The number of different calls used for a given session ranged from 47 to 744. Calls were played for the entire duration of the sessions, with uniformly distributed inter-call intervals. The uniform distribution was from 1.5 to 3.5 s for five sessions, from 0.8 to 1.8 s for two sessions, and from 1 to 2 s for one session. At the end of each inter-call interval, a new call was randomly picked from the set of calls for that session (with replacement) and played. The playback was designed to provide an auditory experience that is similar to the near-constant chatter of bat calls in a bat cave in the wild or in our bat colony room, as was done previously (Prat et al., 2017). The calls used in playback sessions were recorded from the one-chamber sessions. On some of the playback sessions, the calls played to the bats were recorded from the same bats on one-chamber sessions; on other sessions, the calls played were recorded from different bats. Results from neural analyses were similar regardless of the range of inter-call intervals used and the set of calls played, and were thus combined together.

#### 3.2 Behavior definitions and annotations

The behaviors of the bats in our experiments were manually annotated using a custom annotation program written in MATLAB (MathWorks). The annotations were done by experienced trained observers who did not know about the goals of the experiment, the analyses being performed, or the nature of the neural data, and were therefore unbiased. Annotations were done at a detailed level: the behaviors of each bat at each video frame were classified, according to a set of definitions, which we defined after extensive observation of bat behavior. Annotation was done for 65 of the 70 sessions (technical errors in the video recording prevented annotation for the other 5 sessions). It is important to note that due to the fine-grained annotation procedure and the length of the sessions (frame-by-frame annotation of videos recorded at 100 frames/s for sessions lasting ∼100 minutes each), annotation for a single session typically takes between 1-2 months for a single person. Yet, considering that the fine-grained social behavior of this species of bats has not be characterized before, we chose to take this careful, ethological approach despite its time-consuming nature.

Here we detail the definitions of the different behaviors observed in our experiments. They represent the behavioral repertoire the bats exhibited in the experiments, and were defined based on extensive examination of the video recordings of the experiments.

- *Resting*. A bat hanging by its feet, with its head and body still.
- *Active non-social*. A bat engaging in any kind of active behavior that doesn’t involve social interaction, including: the bat hanging by its feet or feet and thumbs, and moving its head or body; the bat climbing or crawling around; the bat shaking its body; the bat jumping or flying off from the roof of the cage.
- *Self-grooming*. A bat either licking or scratching itself.
- *Social grooming*. A bat either licking or scratching another bat.
- *Probing*. A bat poking its snout at the head or body of another bat.
- *Fighting*. A bat moving its wings or thumbs to quickly hit another bat, or biting another bat.
- *Mating*. A male bat inserting or attempting to insert its penis into a female bat’s vagina.
- *Wing covering*. A bat struggling with another bat in order to cover the other bat’s body with its opened wings.
- *Reaching*. A bat attempting to reach over the body or wings of another bat with its head or thumbs.
- *Blocking*. A bat using its wings to actively block another bat from accessing a location.
- *Other interactions*. Any social interaction other than the ones already defined.

#### 3.3 Surgery

Anesthesia and surgical procedures generally followed those described previously in detail for Egyptian fruit bats (Yartsev and Ulanovsky, 2013). Surgeries were performed to implant a four-tetrode microdrive on each bat. Anesthesia was induced using an injectable cocktail of ketamine (22 mg/kgBW), dexamedetomidine (0.09 mg/kgBW) and midazolam (0.31 mg/kgBW). Subsequently, the bat was placed in a stereotaxic apparatus (Kopf) and anesthesia was maintained throughout surgery by repeated injections (roughly once per hour) of an anesthesia maintenance cocktail of dexamedetomidine (0.125 mg/kgBW), midazolam (2.5 mg/kgBW) and fentanyl (0.025 mg/kgBW). The depth of anesthesia was monitored by testing toe pinch reflexes and measuring the bat’s breathing rate. The body temperature of the bat was kept constant at approximately 35-36°C, using a closed-loop temperature controller (FHC) connected to a rectal temperature probe and a heating pad placed under the bat.

Each bat was implanted with a four-tetrode lightweight microdrive (Harlan 4 Drive, Neuralynx; weight 2.1 g). Tetrodes (∼45 µm diameter) were constructed from four strands of platinum-iridium wire (17.8 µm diameter, HML-insulated), bound together by twisting and then melting their insulations. Each of the four tetrodes was loaded and glued separately into a telescoped assembly of polyimide tubes mounted into the microdrive. The tetrodes exited the microdrive through a guide cannula in an approximately rectangular arrangement with ∼300 µm horizontal spacing between tetrodes. Each tetrode could be moved independently via a separate drive screw. On the day before surgery, the tip of each tetrode was cut flat using high-quality scissors (tungsten-carbide scissors with ceramic coating, CeramaCut; FST) and plated with Platinum Black (Neuralynx) to reduce the impedance of individual wires to 0.3-0.8 MΩ (at 1 kHz).

While the bat was under anesthesia, the skull was micro-scarred to improve subsequent adhesion, and a circular opening (craniotomy of 1.8 mm diameter) was made in the skull over the left hemisphere. The center of craniotomy was positioned over the frontal cortex of the bat at 1.7 mm lateral to the midline and 12.19 mm anterior to the transverse sinus that runs between the posterior part of the cortex and the cerebellum. After removal of the dura, the microdrive was lowered and the tip of the microdrive’s guide tube was placed on the brain surface. The microdrive was placed vertically. The craniotomy was then filled with a biocompatible elastomer (Kwik-Sil, World Precision Instruments) to protect the brain. The exposed muscle tissue was then covered with a thin layer of biocompatible adhesive (Vetbond, World Precision Instruments) for protection. A bone screw (FST) with a soldered stainless-steel wire was fixed to the skull in the frontal plate, and served as a ground screw after its electrical connection to the dura was verified. An additional set of 3-5 bone screws were fixed to the skull and served as anchor screws for the mechanical stability of the implant. The bases of the screws were then covered with a thin layer of quick adhesive cement (C&B Metabond, Parkell) which held the screws firmly to the skull; dental acrylic was then added to secure the entire microdrive to the screws and to the skull. At the end of the surgery, bats were given the analgesic Metacam and the anti-inflammatory drug dexamethasone.

#### 3.4 Electrophysiological recording

Electrophysiological recordings were conducted using a wireless neural data logging system (Neurolog-16, Deuteron Technologies), which amplifies the voltage signals from the 16 channels of the 4 tetrodes, performs analog-to-digital conversion at a sampling rate of 29.29 kHz, and stores the digitized data on an on-board SD card. The system has a bandwidth of 1 Hz - 7 kHz, records voltage with a fine resolution of 3.3 µv, and has a low level of noise generally close to the limit of Johnson noise from the impedance of a given source. The system also contains infrared LEDs that can be turned on and off during recording, whose on and off time stamps are recorded along with time stamps of neural data; these LEDs were used to synchronize video and neural recording (see above). Furthermore, the recording system is light-weight (9.9 g, including battery and plastic casing). The Egyptian fruit bats used in our experiment weighed more than 160 g and carried the recording system with ease, as expected from previous experiments using wireless recording systems with heavier or comparable weights during free flight for over an hour and covering multiple kilometers (Yartsev and Ulanovsky, 2013; Finkelstein et al., 2014).

After all recording sessions were concluded for the day, we connected the tetrodes to a wired recording system (Digital Lynx, Neuralynx) to monitor the neural signals and advance the tetrodes. Tetrodes were moved downward once every one to two days (mostly by 20-160 µm), in order to record single units, local spiking activity and LFP at new sites.

#### 3.5 Histology

Histology was done as described previously (Yartsev and Ulanovsky, 2013). Each bat was given a lethal overdose of sodium pentobarbital and, with tetrodes left *in situ*, was perfused transcardially using a flush of 50 ml phosphate-buffered saline followed by 200 ml of fixative (4% paraformaldehyde + 0.1 M phosphate-buffered saline). The brains were then removed and stored in fixative. Subsequently, a cryostat was used to cut 40 µm coronal sections of the brains. The sections were Nissl-stained with cresyl violet, and cover-slipped. A light microscope fitted with a digital camera was used to determine tetrode locations.

#### 3.6 Preprocessing of electrophysiological data

All filtering described in this section were done twice, in the forward and reverse directions, to eliminate phase distortion.

To obtain LFP, we first low-pass filtered the raw voltage traces using a 10th-order Butterworth filter with a cut-off frequency of 200 Hz. The voltage traces were then downsampled by a factor of 59, resulting in a sampling frequency of 496.6 Hz. Power line noise was then filtered out using a 10th-order Butterworth band-stop filter with cut-off frequencies 59.5 Hz and 60.5 Hz, and another one with cut-off frequencies 119.5 Hz and 120.5 Hz.

We observed artifacts in our LFP recording, in the form of large amplitude, transient (∼200 ms), irregular voltage fluctuations that are visually distinct from the normal LFP signal. To automatically detect these artifacts, we used the following algorithm. We note that this algorithm is only a heuristic method that worked well for our data; while it is convoluted, it performed better than a number of simpler methods we tried.

For a given LFP voltage trace (from one recording channel, spanning one recording session), we calculate its spectrogram over our frequency band of interest, 1-150 Hz. Specifically, modified periodograms are computed for short, overlapping segments (64 samples, or ∼128.89 ms, with 50% overlap between consecutive segments) of the LFP trace, each windowed with a Hamming window. In the spectrogram, artifacts appear as spikes in power. To facilitate the detection of these power outliers at any frequency, we normalize the spectrogram as follows: for each frequency, we normalize the power at that frequency by the median absolute deviation of power at that frequency. This normalized spectrogram is a matrix (number of frequencies X number of time bins), which we denote by *S*. We average over the rows of *S* (i.e. averaging across frequencies) to obtain a vector *M* (1 X number of time bins), which is the average normalized power as a function of time. We set a threshold, *M*_*threshold*_, to be the median of *M* multiplied by a parameter *T*_*m*_. Elements of *M* that are larger than *M*_*threshold*_ are detected as potential artifacts. *T*_*m*_ was chosen separately for each recording channel on each session, based on manual inspection of the artifact detection results obtained using a range of *T*_*m*_ values. The median *T*_*m*_ across all recording channels, bats, and sessions (n = 1912) is 8, and the first and third quartiles are 5 and 12, respectively.

The detected potential artifacts could include normal large amplitude oscillations that occur during sleep. To detect these false positives, we used the following procedure. At each time bin that a potential artifact is detected, we take the corresponding column of *S*, and find the maximum in that column between 1 and 10 Hz, which we denote by *P*_*low*_. Similarly, we find *P*_*mid*_ as the maximum between 10 and 20 Hz, and *P*_*high*_ as the maximum between 45 and 120 Hz. Then, we classify a potential artifact as a false positive if all three of the following criteria are satisfied: (1) *P*_*mid*_ / *P*_*high*_ > 6.5; (2) *P*_*mid*_ / *P*_*low*_ > 2; (3) the element of *M* at the given time bin is smaller than 1.5 *M*_*threshold*_.

Each element of *M* corresponds to 64 voltage samples. After rejecting the false positives, for each remaining element of *M* that is larger than *M*_*threshold*_, we define the corresponding 64 voltage samples as a single artifact. For consecutive elements of *M* that are larger than *M*_*threshold*_, their corresponding voltage samples are merged into a single artifact. Then, we define a voltage range within which normal LFP signal lies: the median ± 3 times the median absolute deviation of the entire voltage trace. For each artifact, if the first sample before it or the first sample after it is not within the normal voltage range, then we extend the artifact until both are within the normal range; this makes sure that the algorithm catches the “tails” of each artifact. Then, if the interval between any two artifacts is shorter than 210 ms, the two artifacts and the interval between them are merged into a single artifact.

When analyzing LFP, after we remove an artifact from an LFP trace, we close the resulting gap by joining the two ends of the trace. If the voltages at these two ends differ by a large amount, this effectively creates a new artifact, which we would like to avoid. Thus, before artifact detection, for a given voltage trace, we calculate the absolute value of the voltage difference between every pair of consecutive samples. We define the 90th percentile of all these absolute values as the largest acceptable voltage difference across the two ends of an artifact. If the voltage difference across the two ends of an artifact is larger than this threshold, we extend the artifact by up to 100 ms on each side, to bring the difference below the threshold, making sure to extend by the minimal length possible. If extensions by up to 100 ms on each side are not enough to bring the difference below threshold, we choose the lengths of extensions (still constrained to be below 100 ms on each side) to minimize the difference.

In total, artifacts amounted to a small proportion of our recordings. For each recording channel on each session, we calculated the total duration of artifacts, and the total duration of artifacts as a proportion of the total recording duration. For the total artifact duration, the median across all recording channels, bats, and sessions (n = 1912) was 36.7 s, and the first and third quartiles were 14.1 s and 126.7 s, respectively. For the total duration of artifacts as a proportion of the total recording duration, the median was 0.0058, and the first and third quartiles were 0.0023 and 0.020, respectively. For all the LFP analyses presented in this paper, artifacts were removed prior to analysis, as described in “Calculation of LFP spectrograms” below.

To detect spikes, we band-pass filtered the raw voltage traces using a 6th-order Butterworth filter with cut-off frequencies of 600 Hz and 6000 Hz. For each recording channel and each session, a voltage threshold was set as the following quantity: the difference between the 75th percentile and the median of the voltage trace, divided by the 75th percentile of the standard normal distribution, and multiplied by a factor of 3 (Quian Quiroga et al., 2004). Each time the voltage on one recording channel crossed its threshold, we found the sample having the peak voltage among the over-threshold samples, and extracted 32 samples (1.09 ms) from each channel of the tetrode around the time of the peak sample: from the 7th sample before the peak sample to the 24th sample after. These extracted samples were then used for spike sorting. We performed spike sorting automatically using SNAP Sorter (Neuralynx) with the default settings, then manually checked and cleaned up the results using SpikeSort3D (Neuralynx). For each tetrode on each session, after identifying spikes belonging to single units and after excluding artifacts based on waveform shape, all remaining spikes were grouped into a multiunit. All units with firing rate below 2 Hz were excluded from further analysis. Our dataset included a total of 326 single units and 530 multiunits (after excluding units with low firing rates).

#### 3.7 Calculation of LFP spectrograms

For each LFP trace, we calculated its spectrogram as follows. Power spectra were calculated for 5 s sliding windows of the LFP trace, with 2.5 s overlap between consecutive windows. The window size of 5 s was chosen as the shortest window that resulted in tolerable levels of noise in the spectral estimates, as assessed by visual inspection of the power spectra. The power spectra were computed at integer frequencies from 1 to 150 Hz, using the multitaper method with a time half bandwidth product of 4. If a given window contained artifacts whose duration exceeded 3.5 s, we did not compute a power spectrum for that window, and instead interpolated its power spectrum from those of the neighboring windows. If a given window contained artifacts whose duration did not exceed 3.5 s, we computed its power spectrum after removing the artifacts. Thus, for all the LFP analyses presented in this paper, artifacts were removed prior to analysis. Furthermore, power at 60 Hz was interpolated from power at 59 and 61 Hz, and power at 120 Hz was interpolated from power at 119 and 121 Hz.

To analyze and visualize different frequencies on equal footing, for each LFP spectrogram, we separately peak-normalized the power at each frequency, i.e. power at each frequency was divided by the peak power at that frequency. Other methods of normalization and whitening (such as z-scoring the power at each frequency) gave similar results, so we opted for the simple method of peak-normalization.

#### 3.8 Calculation of firing rates

For single units and multiunits, firing rate as a function of time was calculated in 5 s bins with 2.5 s overlap between consecutive bins. The bin size and overlap were chosen to be the same as for the computation of LFP spectrograms, so that the LFP and spiking results can be compared.

#### 3.9 Dimensionality reduction of LFP

To reduce the dimensionality of the normalized LFP spectrograms, we performed PCA on them, with frequencies as variables and time bins as observations. PCA consistently identified two dimensions that stood out from the noise (Zhang and Yartsev, 2019), which we approximated with flat frequency bands, as they are easier to interpret. The flat frequency bands were chosen as follows. We can divide the range of 1-150 Hz into two frequency bands defined by a dividing frequency, e.g. a dividing frequency of 29 Hz divides the range of 1-150 Hz into the 1-29 Hz band and the 30-150 Hz band. For each normalized spectrogram (corresponding to one recording channel, bat, and session), we calculated the combined variance captured by two flat frequency bands, divided by the combined variance of the top two PCs (which is the maximum amount of variance that can be captured by two dimensions), as a function of the dividing frequency. The dividing frequency at which this variance proportion is maximized is the optimal dividing frequency for this normalized spectrogram. To determine the optimal dividing frequency for all normalized spectrograms (i.e. all channels, bats, and sessions), we averaged the variance proportion curves across all normalized spectrograms: the peak of the averaged curve occurred at 29 Hz. Consistent with this, the median across the optimal dividing frequencies of the different normalized spectrograms was also 29 Hz, and the first and third quartiles were 27 Hz and 32 Hz, respectively. Thus, we chose to reduce the dimensionality of the normalized LFP spectrograms to the 1-29 Hz band and the 30-150 Hz band.

To analyze power over time in each of these frequency bands, for a given normalized LFP spectrogram, we averaged the normalized power values across frequencies within the band. This is the “mean normalized LFP power” used for all analyses in this paper involving LFP power (e.g. Figure 1B).

#### 3.10 Detection of social calls

We used the following procedure to automatically detect social calls from audio recordings of the experimental sessions. A subset of detected calls was used for playback in the two-chambers playback sessions, as described above. Given an audio signal (normalized to the range from -1 to 1, the range of the recording system), its envelope was calculated as the magnitude of its analytic representation (a complex function whose real part is the audio signal and whose imaginary part is the Hilbert transform of the audio signal). The envelope was low-pass filtered in both the forward and reverse directions using a 3rd-order Butterworth filter with a cut-off frequency of 750 Hz. Putative calls were detected as threshold-crossing by the filtered envelop at a threshold of 0.002. Putative calls with a root mean square below 0.01 or a duration shorter than 15 ms were discarded as false positives; all others were detected as calls.

Consecutive calls whose inter-call interval was shorter than 20 ms were merged into a single call.

### 4 ANALYSIS AND MODELING

#### 4.1 Statistical tests

The statistical tests used are stated in the figure legends. A significance level of 0.05 was used for all tests. Tests were two-tailed unless otherwise indicated.

#### 4.2 Inter-brain difference and mean components: definition and quantification

The neural activity of two bats can be represented as a two-dimensional vector 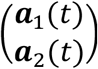, where ***a***_1_(*t*) and ***a***_2_(*t*) are the neural activity (normalized LFP power, multiunit activity, or single unit activity) of bat 1 and bat 2 at time *t*, respectively. Through a change of basis, the same activity can be represented under another orthogonal basis as the mean and difference between the two brains: 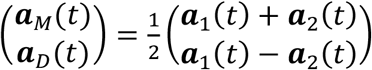, where ***a***_*M*_ (*t*) is the mean component of the activity, and ***a***_*D*_(*t*) is the difference component. Here, the difference component is defined as 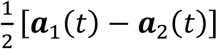 rather than ***a***_1_(*t*) − ***a***_2_(*t*) so as to have the same scale as the mean component. ***a***_*M*_(*t*) represents the common activity pattern shared between the brains, and ***a***_*D*_(*t*) represents the activity difference between the brains.

To quantify the magnitude of an activity component on a given session, we computed the variance of its time series over the session. To quantify the timescales of an activity component on a given session, we computed its power spectral centroid as follows. Given the time series of an activity component over the session, we subtracted from it its average over time, multiplied it with a Hamming window, and then computed the periodogram estimate of its power spectrum. The power spectral centroid is then computed as a weighted average of frequency, with the power at each frequency as its weight. A higher power spectral centroid means that the activity has more power at higher frequencies, so that the time series varied at faster timescales. Note that the power spectral centroid was always calculated from time series of mean normalized LFP power or firing rate, and not from time series of LFP itself.

#### 4.3 Surrogate data and the relationship between inter-brain correlation and mean and difference components

In our data from one-chamber sessions, the mean component had large magnitudes and slow timescales, and the difference component had small magnitudes and fast timescales. The relative magnitudes of the two components are expected, given that activity from the two bats showed similar patterns over time (Figure 1B), with high positive inter-brain correlations (Zhang and Yartsev, 2019); as explained below, a positive correlation mathematically implies larger variance for the mean component compared to the difference component. What about their timescales? Are the observed relative timescales of the two components necessary mathematical consequences of high inter-brain correlations, large mean components, and small difference components?

To answer this, we examine the relationship between inter-brain correlation and the mean and difference components. Let’s use ***A***_1_, a column vector, to denote the activity of bat 1 during a session, and similarly, ***A***_2_ for the activity of bat 2. ***A***_1_ and ***A***_2_ are *N*-dimensional vectors, where *N* is the number of time points in the session. The activity of the mean component is ***A***_*M*_ = (***A***_1_ + ***A***_2_)/2, and the activity of the difference component is ***A***_*D*_ = (***A***_1_ − ***A***_2_)/2. We use ***Â***_1_ to denote the mean-subtracted version of ***A***_1_ (here “mean-subtracted” refers to subtracting the average across the elements of a vector, i.e. across time, not to be confused with the average across bats, as in the “mean component”), and similarly, ***Â***_2_, ***Â***_*M*_, and ***Â***_*D*_ for mean-subtracted versions of the respective vectors. Then, the Pearson correlation coefficient between the activity of two brains is:

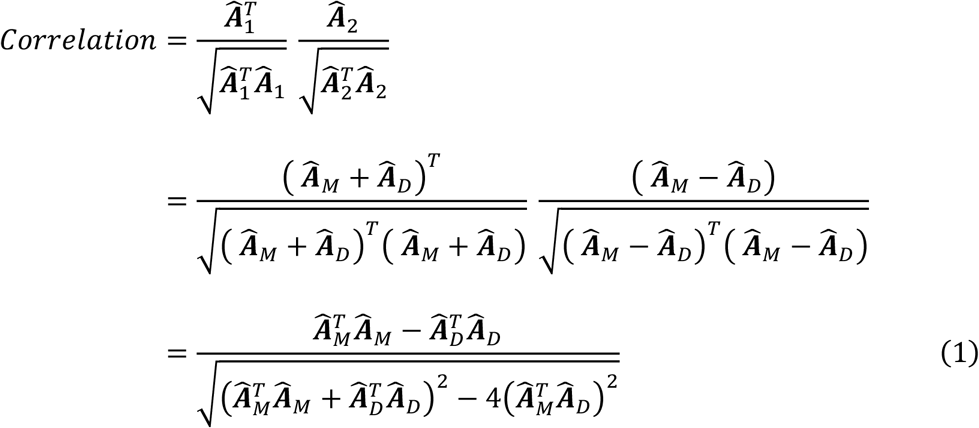

Thus, the correlation depends on the quantities 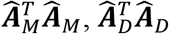, and 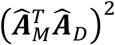. Note that 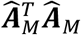 is *N* times the variance of the mean component, and 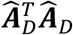 is *N* times the variance of the difference component. A positive correlation requires the numerator in equation (1) to be positive, in other words, the mean component having larger variance than the difference component, as stated above.

Equation (1) also shows that having a given combination of correlation, mean component variance, and difference component variance does not place constraints on the timescales of the mean and difference components. Specifically, it does not constrain the difference component to be faster than the mean. To explicitly demonstrate this, we generated surrogate data with identical inter-brain correlation, mean component variance, and difference component variance as actual data, but having a slower difference component than the mean component (Figure 2).

The surrogate data in Figure 2A-C were generated from the actual 30-150 Hz LFP power data of the example session shown in Figure 1B-D, by keeping the mean component of the actual data, and replacing the difference component with a surrogate, using the following procedure.

Below we denote the actual data by ***A***_*D*_, ***Â***_*D*_, etc. as above, and denote their surrogate counterparts by ***S***_*D*_, *Ŝ*_*D*_, etc. We generated a random *N* × 1 activity vector by picking each element independently from the uniform distribution between 0 and 1, smoothed it with a 1000-second moving average filter, and subtracted from it the average across its elements. Let’s call the resulting vector 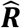. Due to the smoothing, 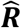 varied on slow timescales. We then found the component of 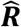 that is orthogonal to 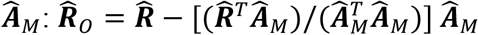. Next, we constructed a vector 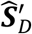 as a linear combination of ***Â***_*M*_ and 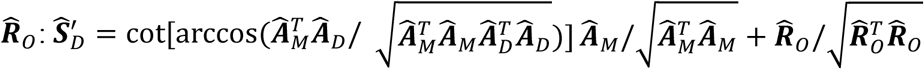. The surrogate for ***Â***_*D*_ is the vector *Ŝ*_*D*_, obtained by scaling 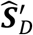 to have the same vector norm as 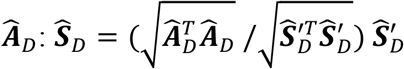. The surrogate difference component is then ***S***_*D*_ = *Ŝ*_*D*_ + ***Ā***_*D*_, where ***Ā***_*D*_ is the average across the elements of ***A***_*D*_. The surrogate activity of bat 1 and bat 2 are, respectively, ***S***_1_ = ***A***_*M*_ + ***S***_*D*_ and ***S***_2_ = ***A***_*M*_ − ***S***_*D*_.This procedure ensures that 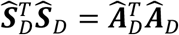 and 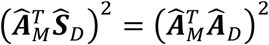, so that the surrogate data had identical inter-brain correlation, mean component variance, and difference component variance as the actual data from which it was generated. Note that here we chose to leave the mean component ***A***_*M*_ from the actual data unchanged in generating the surrogate data, so that the surrogate ***S***_1_ and ***S***_2_ show qualitatively similar behavioral modulation over time as the actual data ***A***_1_ and ***A***_2_, but it’s also possible to replace both ***A***_*M*_ and ***A***_*D*_ with surrogate counterparts.

For Figure 2D-E, we repeated the above procedure for each of the one-chamber sessions from the actual data set, using the actual 30-150 Hz LFP power data.

#### 4.4 Modeling: procedure

Our goal of modeling was to reproduce the observed relationship between difference and mean components using a model that is simple and parsimonious, in order to unambiguously identify the underlying computational mechanisms. We modeled the time evolution of the neural activity of two bats using the following differential equation:

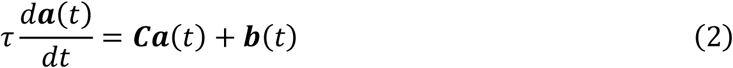

Here *τ* is a time constant; *t* is time; 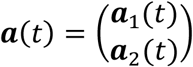 is the activity of bat 1 and bat 2 at time *t*; 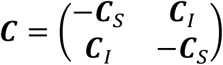 is the functional coupling matrix, where ***C***_*S*_ is the strength of functional self-coupling and ***C***_*I*_ is the strength of functional across-brain coupling; 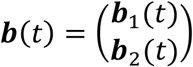 is the strengths of behavioral modulation, where ***b***_1_(*t*) is the modulation of the neural activity of bat 1 by the behavior of bat 1 at time *t*, and similarly for ***b***_2_(*t*).

The functional across-brain coupling term, ***C***_*I*_, represents the indirect influence (as opposed to direct influence from an actual neural connection) one bat’s neural activity has on the other bat’s neural activity. For one-chamber sessions, ***C***_*I*_ > 0, which models positive functional coupling when the two bats share a social environment (Hasson et al., 2012). For example, when bat 1’s neural activity (say, 30-150 Hz LFP power) increases due to its active movements (Gervasoni et al., 2004; McGinley et al., 2015), the movements create sensory inputs to bat 2, which can increase bat 2’s neural activity to the extent that bat 2 is paying attention (Chun et al., 2011; Driver, 2001; Fritz et al., 2007; Reynolds and Chelazzi, 2004). For two-chambers sessions, ***C***_*I*_ = 0 since the two bats are separated. To ensure stability (so that neural activity do not go to infinity), −***C***_*S*_ must be negative and must have a larger absolute value than ***C***_*I*_; thus, 0 ≤ ***C***_*I*_ < ***C***_*S*_.

To generate ***b***(*t*) for a simulation, we first simulated the bats’ behaviors using a Markov chain. Each state of the Markov chain corresponds to the behaviors of the two bats at a given time: e.g. bat 1 resting and bat 2 self-grooming would be one state. For the Markov chain for one-chamber sessions, the transition probability matrix, initial distribution, and state space were determined as follows. The transition probability from one state to another is taken to be the empirical frequency of that transition during all one-chamber sessions. To calculate the empirical frequency, the behavior of each bat was sampled every 2.5 s, at the same time points as the neural activity (i.e. the center time point of each window used to calculate LFP power). A transition was counted for each consecutive pair of time points (in the rare instances where a bat engaged in multiple behaviors at the same time point, one transition was counted for each of the simultaneous behaviors). The initial distribution was taken to be the distribution of the behavioral state at the first time point of each one-chamber session. For these calculations, the two bats were assumed to be symmetrical, in the following sense. Using “AB” to denote the state of “bat 1 engaging in behavior A, bat 2 engaging in behavior B”, we consider the transitions “AB→CD” and “CD→AB” to have the same transition probability. When calculating the empirical transition frequencies, the count for “AB→CD” and the count for “CD→AB” were each taken to be the sum of the actual counts for “AB→CD” and “CD→AB”. This applied also to states where both bats were engaging in the same behavior: e.g., the count for the transition “AA→AA” was taken to be double the actual count. This symmetry assumption was made for simplicity, and to allow behavioral data from different pairs of bats to be pooled together. Once the empirical transition frequencies were calculated, if there were less than 100 transitions from a given state, then that state was excluded from the state space. The same procedures were used to determine the transition probability matrix, initial distribution, and state space for the two-chambers Markov chain. The transition probability matrices for one-chamber and two-chambers sessions are shown in Figure S4A-B.

For each simulated session, we simulated the bats’ behaviors for 100 minutes using the Markov chain (with 2.5 s between steps, the behavioral sampling period used when calculating the empirical transition frequency). The first state of each simulated session was drawn randomly from the appropriate initial distribution. From the simulated behaviors, ***b***(*t*) was determined at the discrete time points of the Markov chain steps (at integer multiples of 2.5 s). At the time point of a given Markov chain step, say *t*_1_, ***b***(*t*_1_) was the sum of the noiseless deterministic behavioral modulation ***b***_*d*_(*t*_1_) and the behavioral modulation noise ***b***_*n*_(*t*_1_). ***b***_*d*_(*t*_1_) depended only on the behaviors of the two bats at time *t*_1_. For example, if bat 1 was resting and bat 2 was engaging in self-grooming at time *t*_1_, then 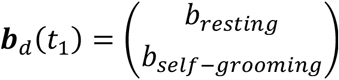, where *b*_*resting*_ and *b*_*self*−*grooming*_ are parameters specifying the level of behavioral modulation associated with the two behaviors. See section 4.10 below for the parameters for all the behaviors. For the behavioral modulation noise, the two elements (for the two bats) of ***b***_*n*_(*t*_1_) was each drawn independently from a Gaussian distribution with zero mean and standard deviation *σ*_*n*_. After determining ***b***(*t*) = ***b***_*d*_(*t*) + ***b***_*n*_(*t*) at the discrete time points of the Markov chain steps, ***b***(*t*) at time points in-between were linearly interpolated.

Having simulated the behaviors and generated ***b***(*t*) for a 100-minute session, we set the initial condition ***a***(0) to be the fixed point under the initial noiseless behavioral modulation ***b***_*d*_(0): ***a***(0) = −***C***^−1^***b***_*d*_(0). Then, equation (2) was numerically integrated (*ode45* function in MATLAB) to simulate neural activity ***a***(*t*). The only differences between simulations of one-chamber and two-chambers sessions were the value of ***C***_*I*_ and the behavior transition probability matrix. For analyses and figures involving ***a***(*t*), ***b***(*t*), ***b***_*d*_(*t*), and ***b***_*n*_(*t*) (Figures 3-6, S4-S6), we used their values taken at discrete time points corresponding to the Markov chain steps (i.e. same sampling period as the data).

#### 4.5 Modeling: magnitudes and timescales of mean and difference components

How is the model able to reproduce the set of experimental observations relating the magnitudes and timescales of the mean and difference components (Figure 3E-F)? To understand the mechanisms in the model, we examine equation (2). In general, in equations of this form, the activity variables are coupled: for example, in the one-chamber model, the activity of bat 1 ***a***_1_(*t*) influences the activity of bat 2 ***a***_2_(*t*), which in turn feeds back on ***a***_1_(*t*). It is easier to understand how the activity evolves if we express the equation in a basis in which the activity variables uncouple. This basis consists of the eigenvectors of the functional coupling matrix ***C***.The eigenvectors of ***C*** are 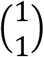 and 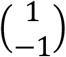, with respective eigenvalues −***C***_*S*_ + ***C***_*I*_ and −***C***_*S*_ − ***C***_*I*_. We define ***V*** to be the matrix whose columns are the eigenvectors: 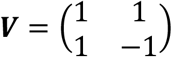. Expressed in the eigenvector basis, the activity variables become the mean activity across bats, and the difference in activity between bats: 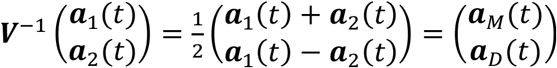. Thus, the uncoupled activity variables are precisely what we are interested in: the mean and difference components. Similarly, in the eigenvector basis, the behavioral modulation variables are the mean behavioral modulation and the difference in behavioral modulation: 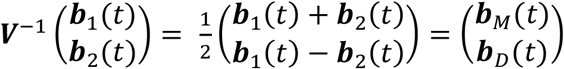.Thus, in the eigenvector basis, equation (2) is

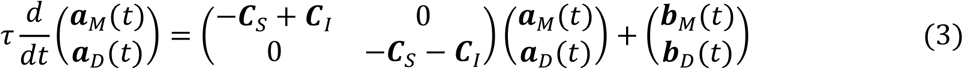

In the one-chamber model, ***C***_*I*_ > 0, so functional across-brain coupling acts as positive feedback to the mean component and negative feedback to the difference component. The effects of such opposite feedback to the two components can be seen more clearly by rewriting equation (3) as the following uncoupled differential equations:

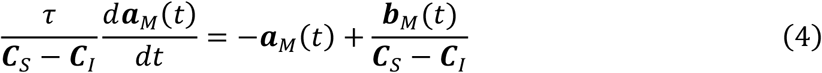

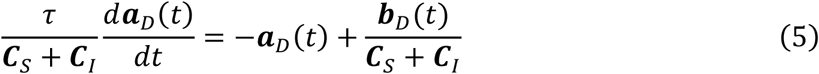

Thus, at any given time *t*, ***a***_*M*_ (*t*) is exponentially approaching 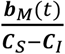 with effective time constant 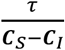, and ***a***_*D*_(*t*) is exponentially approaching 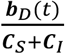 with effective time constant 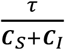. In the one-chamber model, 0 < ***C***_*I*_ < ***C***_*S*_, so 0 < ***C***_*S*_ − ***C***_*I*_ < ***C***_*S*_ + ***C***_*I*_. This means that in the one-chamber model, relative to the two-chambers model where ***C***_*I*_ = 0, the positive feedback provided by functional across-brain coupling amplifies the mean component and slows it down. On the other hand, the negative feedback suppresses the difference component and speeds it up. Thus, the model suggests this opposite feedback to be a potential computational mechanism that explains all of our observations relating the magnitudes and timescales of the mean and difference components.

The only differences between the one-chamber and two-chambers models were the value of ***C***_*I*_ and the behavioral transition probability matrix. Having understood the effects of ***C***_*I*_, we now turn to the effects of the different transition probability matrices. Figure S4C-D shows that, for both the one-chamber and two-chambers models, the respective Markov chains approach stationary distributions in ∼5 minutes, i.e. the distribution of states no longer changes with time. The stationary distribution for the two-chambers model shows that states where the two bats engage in the same behavior are about equally likely as states where they engage in different behaviors (Figure S4E), as expected given that the two bats behave independently in separate chambers. For the one-chamber model, same-behavior states (probability 0.58) are more likely than different-behavior states (Figure S4E). The noiseless behavioral modulation ***b***_*d*_(*t*) depends on the behaviors of the two bats, while the behavioral modulation noise ***b***_*n*_(*t*) does not. Through ***b***_*d*_(*t*), same-behavior states contribute to behavioral modulation of the mean component, and the only noiseless behavioral modulation of the difference component comes from different-behavior states. In the one-chamber model, with increased coordinated behavioral modulation compared to the two-chambers model, the mean noiseless behavioral modulation across bats has larger magnitudes and slower timescales than its difference between bats (blue lines in Figure S4F-G). On the other hand, because the behavioral modulation noise is independent across bats, it does not distinguish between the mean and difference components. Thus, adding the noise to the noiseless behavioral modulation decreases the differential modulation of the mean and difference components (red lines in Figure S4F-G). However, even if the noise strongly reduces differential behavioral modulation of the mean and difference components, the experimentally observed levels of relative magnitudes and timescales can still result from functional across-brain coupling (purple lines in Figure S4F-G).

Thus, our model suggests two mechanisms behind our observations on the mean and difference components: opposite feedback by functional across-brain coupling, and coordinated behavioral modulation. Due to these mechanisms, in a shared social environment, the activity of the two bats become dominated by their common activity pattern, and when the activity of the two bats diverge, they rapidly catch up to each other. As a result, the activity of the two socially interacting bats are highly correlated with each other. We found that inter-brain correlation remained even when the bats were engaged in different behaviors (Figure S5A-B; see also Zhang and Yartsev, 2019), which suggests that coordinated behavioral modulation alone is not sufficient to explain the data, and that opposite feedback by functional across-brain coupling is needed. To explicitly test this, we performed an additional set of simulations of the one-chamber model. Here, after we generated each instantiation of ***b***(*t*), we used it to simulate neural activity twice: once with functional across-brain coupling and once without. Note that the same ***b***(*t*) is used for both simulations, including the same ***b***_*d*_(*t*) generated from the one-chamber Markov chain and the same instantiation of randomly generated ***b***_*n*_(*t*). Figure S5C, D, G shows that the model with functional across-brain coupling reproduced the experimental observation: inter-brain correlation remained after removing periods of coordinated behaviors. On the other hand, without functional across-brain coupling, inter-brain correlation disappeared after removing coordinated behaviors (Figure S5E-G). Thus, we conclude that, in the framework of our model, opposite feedback by functional across-brain coupling is necessary to reproduce the data.

#### 4.6 Modeling: Kuramoto model

We explored an alternative mechanism for inter-brain coupling using the Kuramoto model (Strogatz, 2000; Acebrón et al., 2005). In this model, the activity of the two brains are treated as two oscillators, whose phases are coupled:

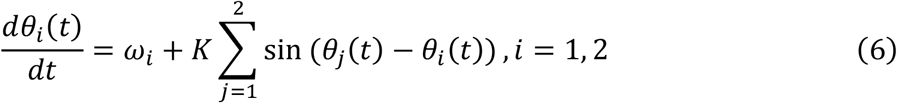

Here *θ*_1_(*t*) and *θ*_2_(*t*) are the phases of the two oscillators at time *t*, and the corresponding activity of the two brains at time *t* are 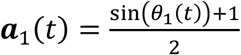and 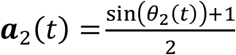; *ω*_1_ and *ω*_2_ are the natural frequencies of the two oscillators; *K* is the coupling strength between the oscillators: *K* > 0 for simulations of one-chamber sessions, and *K* = 0 for simulations of two-chambers sessions.

For each simulation, *ω*_1_ and *ω*_2_ were drawn from lognormal distributions with means 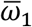 and 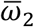, respectively, and the same standard deviation *σ*_*ω*_. 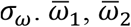, and *σ*_*ω*_ were set to make *ω*_1_ and *ω*_2_ different from each other, to avoid the two brains trivially synchronizing by having the same natural frequencies. To simulate the Kuramoto model, equation (6) was numerically integrated (*ode15s* function in MATLAB) with initial condition *θ*_1_(0) = *θ*_2_(0) = 0. The phases *θ*_1_(*t*) and *θ*_2_(*t*) were then converted to activity ***a***_1_(*t*) and ***a***_2_(*t*), which were plotted in Figure S5H-K and analyzed in Figure S5L-M. This shows that the phase-coupling mechanism of the Kuramoto model is able to reproduce inter-brain correlation and the relative magnitudes of the mean and difference components from the data; however, it does not reproduce the relative timescales of the mean and difference components from the data.

#### 4.7 Modeling: correlation between activity variables under rotated activity bases

The neural dynamics in the model are governed by (2). To express (2) under a different basis, ***a***(*t*), ***C***, and ***b***(*t*) are transformed to ***a***′(*t*) = ***U***^−1^***a***(*t*), ***C***′ = ***U***^−1^***CU***, and ***b***′(*t*) = ***U***^−1^***b***(*t*), respectively, where the columns of the matrix ***U*** are the new basis vectors. If 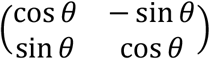, then the new basis corresponds to a counter-clockwise rotation of the original basis (where each axis is the activity of one bat) by an angle of *θ*. By rotating the basis, the functional coupling between the activity variables (i.e. between the two elements of ***a***′(*t*)) changes as a function of the rotation angle (Figure 4A, D). This functional coupling in turn determines the correlations between those activity variables, which forms a set of predictions of the model (Figure 4B, E). These predictions are confirmed in the data in Figure 4C, F.

Additionally, we examined the influence of behavioral modulation on correlation between the activity variables under rotated bases, by regressing out behavior from the activity variables. To do this, for each behavior (e.g. resting, self-grooming, probing, etc.), each bat, and each simulation or experimental session, we used a binary vector to represent the time course of the bat engaging in that behavior: the elements of the vector correspond to time points, and the values of the elements are “1” at time points when the bat was engaging in that behavior, and “0” when it was not. Then, we performed linear regression to predict each activity variable over time under each basis using the set of all behavioral binary vectors of both bats as predictors. The linear regression fit was then subtracted from the activity, and correlation was calculated between these residuals. The results are shown in Figure 4B, C, E, F (brown lines): for both model and data, regressing out behaviors changes the magnitudes, but not the shapes of the correlation curves.

#### 4.8 Modeling: relationship between variance ratio and power spectral centroid ratio

Here we analyze the relationship between the mean/difference ratio of the variance (*r*_*Var*_) and the mean/difference ratio of the power spectral centroid (*r*_*PSC*_) in the model (seen in Figure 4G). Note that, *r*_*Var*_ is not a function of *r*_*PSC*_, and *r*_*PSC*_ is not a function of *r*_*Var*_, because different activity ***a***(*t*) can have different *r*_*Var*_ but the same *r*_*PSC*_, or different *r*_*PSC*_ and the same *r*_*Var*_. But *r*_*Var*_ and *r*_*PSC*_ are both functions of the behavioral modulation ***b***(*t*). Figure 4G shows that *r*_*Var*_ and *r*_*PSC*_ corresponding to different instances of ***b***(*t*) tend to covary linearly. To examine whether this linear relationship is due to a systematic relationship between different instances of ***b***(*t*), or due to the neural dynamics, we examine the effect of random perturbations to behavioral modulation on the relationship between *r*_*Var*_ and *r*_*PSC*_. To do so, we first express *r*_*Var*_ and *r*_*PSC*_ in the frequency domain, and then examine whether and how they covary given random perturbations.

We start with the solutions to (4) and (5), which are, respectively:

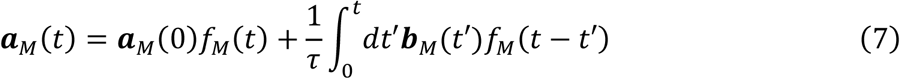

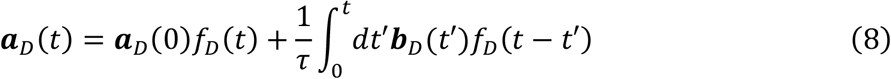

where *f*_*M*_(*t*) and *f*_*D*_(*t*) are the neural decay functions for the mean and difference components:

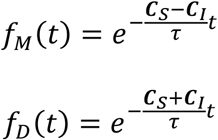

The convolution theorem can be used to approximate the solutions (7) and (8) in the frequency domain:

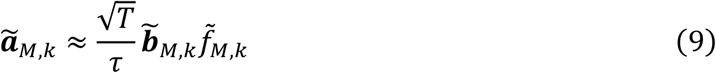

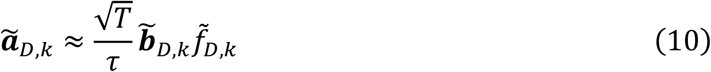

Here, *T* is the duration of the simulated session, and “^∼^” denotes the Fourier transform, e.g., 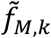 is the Fourier transform of *f*_*M*_(*t*) at frequency 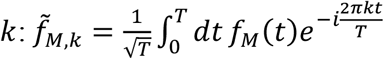. The approximation of (9) and (10) assumes periodic convolutions and ***a***(0) = 0 in (7) and (8), which are not true; nonetheless, the approximation is accurate because the effective time constants 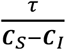 and 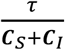 are small compared to *T* in our model.

Using Parseval’s theorem and (9), the variance of the mean component can be approximated as

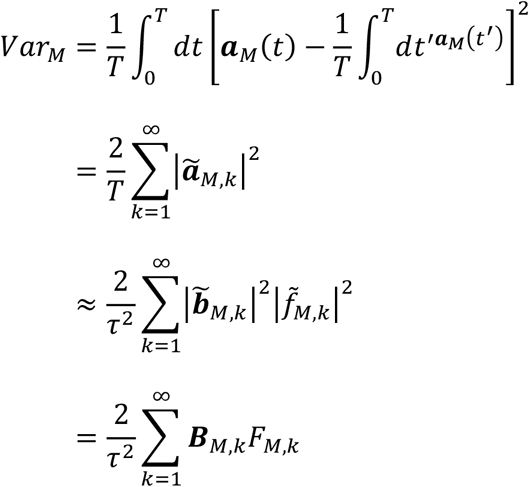

where 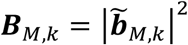 and 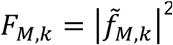. Using (9), the power spectral centroid of the mean component can be approximated as

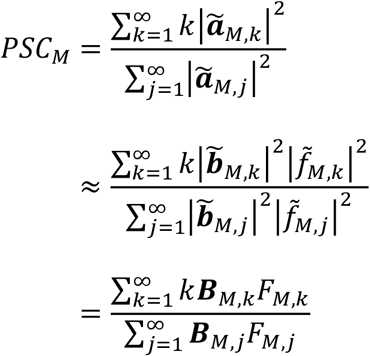

The expressions for the variance and power spectral centroid of the difference component are similar.

Then, the mean/difference ratio of the variance is

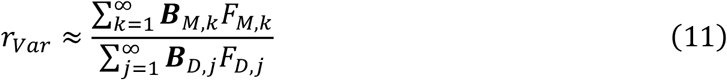

The mean/difference ratio of the power spectral centroid is

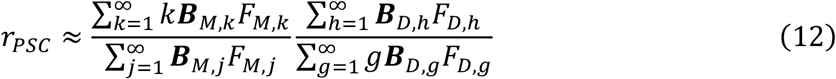

To examine how *r*_*Var*_ and *r*_*PSC*_ change with changes in behavioral modulation, we calculate the following partial derivatives. The partial derivative of *r*_*Var*_ with respect to the power of the mean component of behavioral modulation at frequency *k* is

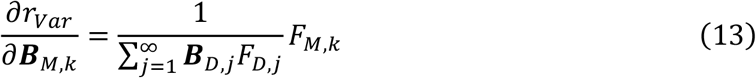

The partial derivative of *r*_*Var*_ with respect to the power of the difference component of behavioral modulation at frequency *k* is

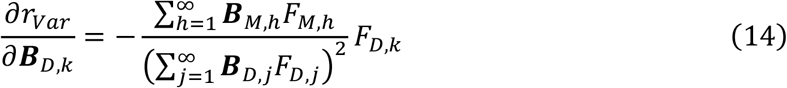

The partial derivative of *r*_*PSC*_ with respect to the power of the mean component of behavioral modulation at frequency *k* is

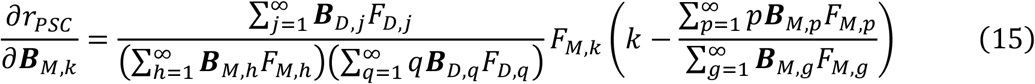

The partial derivative of *r*_*PSC*_ with respect to the power of the difference component of behavioral modulation at frequency *k* is

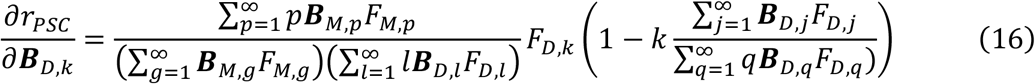

For all calculations here, the infinite sums were truncated at *k*_*t*_ such that 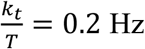, which is the Nyquist frequency for the sampled simulated activity used for our analyses.

We now consider perturbations to the power spectra of behavioral modulation. We concatenate the power spectra of the behavioral modulation of the mean and difference components to form

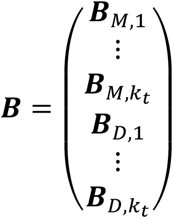

We perturb a given ***B*** by adding *δ****B***, drawn from a uniform distribution on the hypersphere centered at 0. To see whether and how *r*_*Var*_ and *r*_*PSC*_ covary given random perturbations to behavioral modulation, we next determine the covariance matrix between the changes in *r*_*Var*_ and *r*_*PSC*_ resulting from perturbations *δ****B***.

Considering *r*_*Var*_ and *r*_*PSC*_ as functions of ***B***, their gradients are

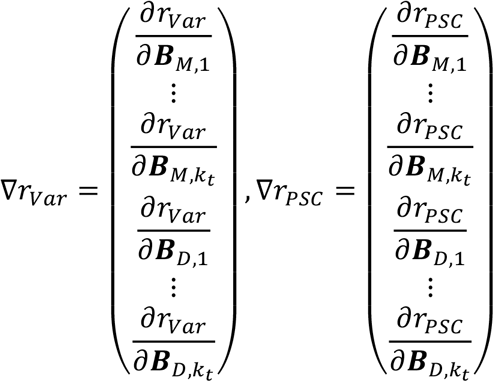

where the partial derivatives are given in (13)-(16). Given a small perturbation *δ****B***, the resulting changes to *r*_*Var*_ and *r*_*PSC*_ are approximately *δ****B***^*T*^∇*r*_*Var*_ and *δ****B***^*T*^∇*r*_*PSC*_, respectively. To calculate the covariance matrix between *δ****B***^*T*^∇*r*_*Var*_ and *δ****B***^*T*^∇*r*_*PSC*_, we first perform an orthogonal transformation to any orthonormal basis whose first basis vector is ∇*r*_*Var*_/‖∇*r*_*Var*_‖, where ‖∙‖ denotes vector norm. We use 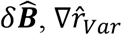, and 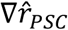 to denote *δ****B***, ∇*r*_*Var*_, and ∇*r*_*PSC*_ under the new basis, respectively. The variance of *δ****B***^*T*^∇*r*_*Var*_ is

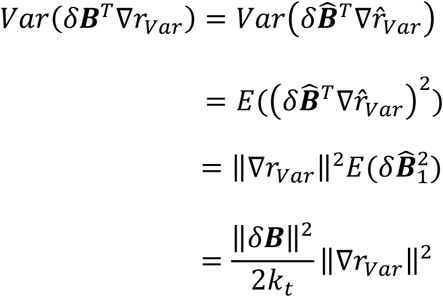

Similarly, the variance of *δ****B***^*T*^∇*r* is 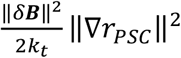. The covariance between *δ****B***^*T*^∇*r*_*var*_ and *δ****B***^*T*^∇*r*_*PSC*_ is

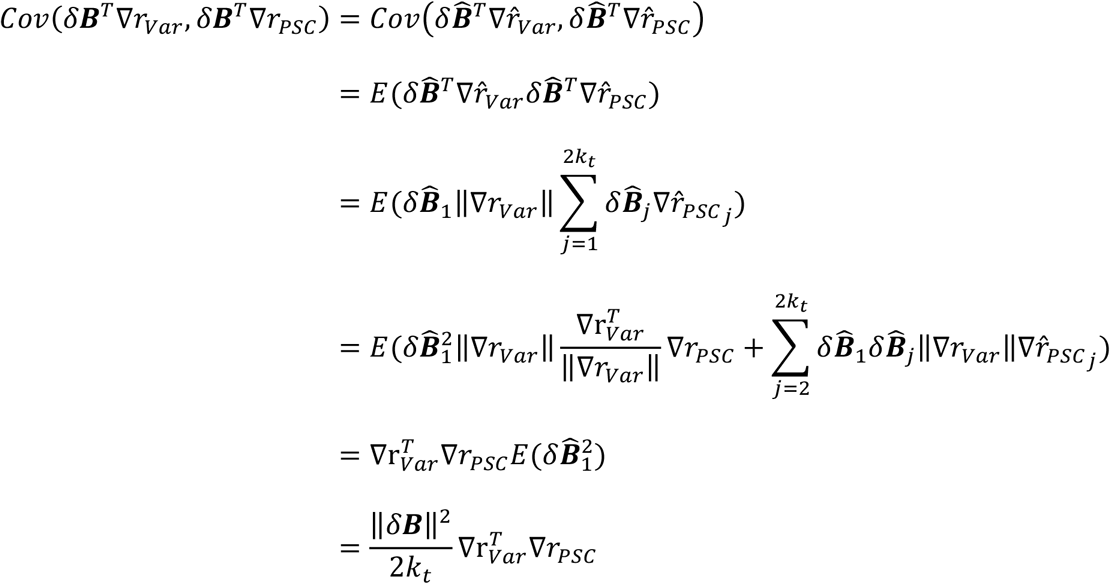

Thus, the covariance matrix between *δ****B***^*T*^∇*r*_*var*_ and *δ****B***^*T*^∇*r*_*PSC*_ is

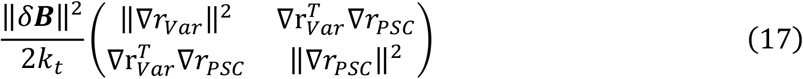

The eigenvector of this matrix corresponding to the larger eigenvalue is the direction of the local linear trend, whereas the relative sizes of the eigenvalues indicate the strength of the linear trend (the larger the difference between the eigenvalues, the stronger the linear trend).

Examining the eigenvectors and eigenvalues of (17) across different simulations (Figure S6; in computing (17), integrals were evaluated numerically), we found that *r*_*Var*_ and *r*_*PSC*_ resulting from random perturbations to ***B*** consistently show linear relationships. Furthermore, the slopes of these local linear relationships, which are not influenced by systematic variations of behavioral modulation ***b***(*t*) across simulations, are consistent with the slopes of the global linear relationships seen in Figure 4G.

#### 4.9 Modeling: using the neural activity of one bat to discriminate the behavior of the other bat

In the model, the neural activity of each bat is directly modulated by its own behavior (e.g., ***a***_1_(*t*) is modulated by ***b***_1_(*t*)). Additionally, in the one-chamber model, the activity of each bat is indirectly modulated by the behavior of the other bat, through functional across-brain coupling (e.g., ***a***_1_(*t*) is modulated by ***b***_2_(*t*) through the coupling ***C***_*I*_). This naturally suggests that the neural activity of each bat should encode the behavior of the other bat independently from encoding its own behavior.

We used the following method to quantify this. Before examining whether the activity of one bat can discriminate the behavior of the other bat, we first regressed out the behavior of each bat from its own neural activity, using the method of section 4.7 above. Importantly, this eliminates potential spurious correlation between one bat’s activity and another bat’s behavior caused by any coordinated behaviors between the bats. We then asked whether the neural activity of a given bat discriminates between a given pair of behaviors by the other bat (e.g., using bat 1’s neural activity to discriminate whether bat 2 is resting or engaging in social grooming), by plotting the receiver operating characteristic (ROC) curve and calculating the area under the curve (Figure 5B, E; Dayan and Abbott, 2005). For each discrimination, positive and negative class assignments for the two behaviors were made so that the area under the ROC curve is greater than 0.5. We then averaged the area under the ROC curve across bats, pairs of behaviors, and simulations or sessions (Figure 5C, F). This showed that the activity of each bat was modulated by the behaviors of the other bat independently of its own behavior, in both the model and the data.

#### 4.10 Modeling: inter-brain relationship during group social interactions

To generalize our two-bat model to more than two bats, we used the same equation (2), with ***a***(*t*) and ***b***(*t*) now being *n*-dimensional vectors, and ***C*** being an *n* × *n* matrix, where *n* is the number of interacting bats. ***C*** retains the same structure from the two-bat model: all diagonal elements (functional self-coupling) are −***C***_*S*_, and all off-diagonal elements (functional across-brain coupling) are ***C***_*I*_.

To understand the *n*-bat model, we examine the eigenvectors and eigenvalues of ***C***. Note that the *n*-dimensional vector whose elements are all 1s is an eigenvector, with eigenvalue (*n* − 1)***C***_*I*_−***C***_*S*_. This eigenvector corresponds to the direction of the mean activity across all bats, and is thus the *n*-bat analogue of the mean component from the two-bat model. Any vector orthogonal to the *n*-bat mean component is also an eigenvector, with eigenvalue −***C***_*I*_−***C***_*S*_. These eigenvectors define an (*n* − 1)-dimensional subspace, which contains all inter-brain activity patterns that correspond to activity differences across brains; we call this subspace the difference subspace, which is the multidimensional analogue of the difference component from the two-bat model. To ensure stability (so that neural activity do not go to infinity), we take our parameter regime to be 0 < (*n* − 1)***C***_*I*_ < ***C***_*S*_. Thus, −***C***_*I*_ − ***C***_*S*_ < (*n* − 1)***C***_*I*_ − ***C***_*S*_ < 0. This means that, similar to the two-bat model, the *n*-bat mean component is amplified and slowed down by the positive feedback provided by functional across-brain coupling, whereas activity patterns in the difference subspace are suppressed and sped up by negative feedback. This results in the predictions that, for *n* socially interacting bats, activity in the *n*-bat mean component will have larger magnitude and slower timescales than activity in the difference subspace on average (Figure 6C-D).

Another prediction concerns the correlation between activity variables during group interactions. Because the activity of pairs of bats are positively functionally coupled in the *n*-bat model, the model would predict positive inter-brain correlations (Figure 6E). On the other hand, because the *n*-bat mean component and vectors in the difference subspace are all eigenvectors, they are functionally uncoupled. Thus, the model would predict zero average correlation between the *n*-bat mean component and activity variables in the difference subspace (Figure 6E).

For the simulations in Figure 6C-E, we used an *n*-bat model with *n* = 4 and performed the simulations using the same procedures as for the two-bat model. Because we do not have behavioral statistics for four-bat group interactions, we opted to use noise for ***b***(*t*), so that model predictions based on functional across-brain coupling would not be biased by structures in ***b***(*t*) (e.g. the *n*-bat mean component could have larger magnitudes than the difference subspace if ***b***(*t*) has the same structure). To generate ***b***(*t*) for the *n*-bat model, we used the following procedure. For a given simulated session and a given bat *i*, we generated ***b***_*i*_(*t*) independently of the other bats. We generated a random vector ***b***_*pre*_, whose dimensionality was the number of time points spaced 2.5 s apart in the simulated session (matching the time step size of the Markov chain from the two-bat model). The elements of ***b***_*pre*_ were drawn independently from a Gaussian distribution with mean *b*_*mean*_ and standard deviation *b*_*std*_. We smoothed ***b***_*pre*_ with a 1200-point moving average filter, then added independent Gaussian noise to its elements (0 mean, standard deviation *σ*_*n*_) as in the two-bat model. The resulting vector contained the values of ***b***_*i*_(*t*) at 2.5 s intervals; ***b***_*i*_(*t*) at time points in-between were linearly interpolated. The same procedure was repeated for each bat.

#### 4.11 Modeling: parameters

In our data, all four neural signals (30-150 Hz LFP power, 1-29 Hz LFP power, multiunits, and single units) show the same qualitative phenomena: faster and smaller difference component compared to the mean component, on one-chamber sessions but not two-chambers sessions (Figure 1). The four signals differed quantitatively in the extent they showed the same qualitative phenomena. The goal of our model is not to quantitatively fit any one of the four neural signals in particular, but to provide mechanistic explanations for the general qualitative phenomena that do not require fine-tuning of parameters. As explained above, the opposite feedback mechanism depends simply on 0 < ***C***_*I*_ < ***C***_*S*_, whereas the coordinated behavioral modulation mechanism is a manifestation of the empirical behavioral transition frequencies.

The focus of the model is to reproduce and explain the qualitative trend of the difference component being faster and smaller than the mean component. Other aspects of the data could be reproduced by extending the model with additional parameters, but we chose not to do so in the interest of keeping the model simple and focused. For example, the 30-150 Hz LFP power data showed more variability across sessions compared to the model simulations (Figure 3E-F). The higher variability could be reproduced if we introduced variability across simulations in the behavioral transition probabilities or the strength of functional across-brain coupling.

The model parameters for the two-bat models are: ***C***_*S*_ = 1, ***C***_*I*_ = 0.4 for simulations with functional across-brain coupling, ***C***_*I*_ = 0 for simulations without functional across-brain coupling, *τ* = 15 s, *σ*_*n*_ = 0.15, *T* = 100 minutes, *b*_*resting*_ = 0.158, *b*_*active non*−*social*_ = 0.269, *b*_*self*−*grooming*_ = 0.264, *b*_*social grooming*_ = 0.223, *b*_*probing*_ = 0.284, *b*_*fighting*_ = 0.355, *b*_*mating*_ = 0.367, *b*_*wing covering*_ = 0.364, *b*_*reaching*_ = 0.321, *b*_*blocking*_ = 0.339, *b*_*other interstions*_ = 0.291. The behavioral modulation parameter for each behavior listed above (*b*_*resting*_, etc.) was set as the average 30-150 Hz mean normalized LFP power during that behavior from the data: take a given bat, for each session, average its 30-150 Hz mean normalized LFP power across all channels, pool together these averaged power values from all time points when the bat engaged in the given behavior from all sessions (including both one-chamber and two-chambers), then pool across all bats, and then average across all such pooled data.

The model parameters for the four-bat model are: ***C***_*S*_ = 1, ***C***_*I*_ = 0.1, *b*_*mean*_ = 0.2, and *b*_*std*_ = 3.5. All other parameters are the same as in the two-bat model.

The model parameters for the Kuramoto model are: *K* = 0.0035 for simulations of one-chamber sessions, *K* = 0 for simulations of two-chambers sessions, 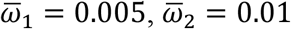, and *σ*_*ω*_ = 0.0005.

#### 4.12 Sample sizes

In this section we list the sample sizes for all results that were presented as averages.

Figure 1M, N, Q, R: n = 52 sessions for one-chamber sessions, and 18 sessions for two-chambers sessions

Figure 1O, S: n = 675 multiunit pairs for one-chamber sessions, and 284 multiunit pairs for two-chambers sessions

Figure 1P, T: n = 256 single unit pairs for one-chamber sessions, and 65 single unit pairs for two-chambers sessions

Figure 3E-F: for data, n = 52 sessions for one-chamber sessions, and 18 sessions for two-chambers sessions; for model, n = 100 simulations for one-chamber model, and 100 simulations for two-chambers model

Figure 4B: n = 100 simulations

Figure 4C: n = 52 sessions

Figure 4E: n = 100 simulations

Figure 4F: n = 18 sessions

Figure 5C: n = 8235 simulations × bats × behavior pairs for one-chamber simulations, n = 1848 simulations × bats × behavior pairs for two-chambers simulations

Figure 5F: n = 2086 sessions × bats × behavior pairs for one-chamber sessions, n = 200 sessions × bats × behavior pairs for two-chambers sessions

Figure 6C-E: n = 100 simulations

Figure S4F, G: for data, n = 52 sessions for one-chamber sessions, and 18 sessions for two-chambers sessions; for model, n = 100 simulations for one-chamber model, and 100 simulations for two-chambers model

Figure S5G: for data, n = 50 sessions; for model, n = 100 simulations with functional across-brain coupling, and 100 simulations without functional across-brain coupling

Figure S5L-M: for data, n = 52 sessions for one-chamber sessions, and 18 sessions for two-chambers sessions; for the Kuramoto model, n = 100 simulations for one-chamber model, and 100 simulations for two-chambers model

Figure S6C-E: n = 100 simulations for one-chamber model, and 100 simulations for two-chambers model

